# Lineage specific transcription factor waves reprogram neuroblastoma from self-renewal to differentiation

**DOI:** 10.1101/2020.07.23.218503

**Authors:** Deblina Banerjee, Berkley Gryder, Sukriti Bagchi, Zhihui Liu, Hsien-Chao Chen, Man Xu, Ming Sun, Zalman Vaksman, Sharon J. Diskin, Javed Khan, Carol J Thiele

## Abstract

Temporal regulation of super-enhancer (SE) driven lineage specific transcription factors (TFs) underlies normal developmental programs. Neuroblastoma (NB) arises from an inability of sympathoadrenal progenitors to exit a self-renewal program and terminally differentiate. To identify critical SEs driving TF regulators of NB, we utilized NB cells in which all-trans retinoic acid (ATRA) induces growth arrest and differentiation. H3K27ac ChIP-seq paired with RNA-seq over a time course of ATRA treatment revealed SEs moving in a coordinated manner with four distinct temporal patterns (clusters). SEs that decreased with ATRA linked to 24 TFs involved in stem cell development/specialization (*MYCN, GATA3, SOX11)* along with *LMO1*, a transcriptional coregulator and oncogene identified via a genome-wide association study (GWAS) of NB. H3K27ac levels and GATA3 binding at the NB-associated rs2168101 site of the LMO1 SE were reduced with ATRA treatment, resulting in 1.46 fold decreased *LMO1* expression. The SOX11 SE was lost coincident with a 50% decrease in mRNA after 8 days of ATRA treatment. CRISPR-Cas9 screening and siRNA inhibition showed a dependency on SOX11 for cell growth in NB cell lines. Silencing of the SOX11 SE using dCAS9-KRAB targeted guides caused a 40% decrease in *SOX11* mRNA and inhibited cell growth. Three other TF SE clusters had sequential waves of activation at 2, 4 and 8 days of ATRA treatment and involved TFs regulating neural development (*GATA2* and *SOX4*). Silencing of the gained SOX4 SE using dCAS9-KRAB targeting, caused a 50% decrease in SOX4 expression and attenuated expression of ATRA-induced differentiation genes. Our study has identified candidate oncogenic lineage drivers of NB self-renewal and TFs critical for implementing a differentiation program.

## Introduction

Neuroblastoma is a tumor of the peripheral sympathetic nervous system, where despite intensive multimodal therapies, survival is limited to 50% in high-risk cases^1^. Patients with undifferentiated or stroma poor tumors have a poor prognoses and tumors overexpress genes regulating cell cycle progression and may have MYCN amplification or copy number alterations (1p36del; unbalanced 11q; 17q+; 7+). In contrast, low risk patients have mature, stroma-rich tumors (ganglioneuromas) with a range of differentiated phenotypes and a transcriptome enriched in differentiation genes^2-4^. High-risk neuroblastoma tumors are believed to be a disease resulting from a differentiation block and may represent a transitional epigenetic state. This has led to several studies describing the super-enhancer (SE) mediated transcriptional core regulatory circuits (CRCs) governing gene expression programs that drive neuroblastoma growth and survival^5-7^. Several lines of investigation indicate that genetic alterations in NB are associated with oncogenic subversion of the transcription factors normally involved in neural crest development^8^. However, how this regulatory circuitry is perturbed when tumor cells switch from a self-renewal state to a differentiated state has been relatively unexplored. Understanding the molecular mechanisms regulating these processes may reveal insights that can be therapeutically exploited.

Current models indicate NB originates from a failure of neural crest derived sympathoadrenal progenitor cells to exit a self-renewal program and implement a differentiation program^9^. At a molecular level, lineage specification and differentiation are complex processes dictated by gene regulatory networks which are controlled by a small number of master transcription factors (MTFs)^10^ which are often involved in CRCs. Cancer cells often inherit their transcriptional profile from their normal cellular counterparts. Their abnormal growth may involve lineage subversion rendering cells unable to respond to physiologic cues that normally regulate their growth and differentiation programs. Thus, one current approach to understand cancer and identify critical therapeutic targets, is to delineate key transcriptional machinery dependencies^11^. The lineage-dependency mechanisms that promote tumor progression often involve lineage master regulatory genes. A recent study has shown that several master regulators important in neural crest lineages including PHOX2B, HAND2, form a CRC on which the neuroblastoma tumor cells are dependent for survival and growth^6^. Thus, like lineage regulating MTFs, they are expressed at high levels and are driven by cell type specific super-enhancers^11^.

During development, trunk neural crest cells delaminate from the neural tube and undergo neural differentiation in which some commit to the sympathoadrenal lineage^12^. MYCN, an oncogene which plays a role in aggressive NB, is expressed throughout the process of migration, survival and/or differentiation of these migrating neural crest cells and its levels gradually reduce in differentiating sympathetic neurons^13,14^. However, in neuroblastoma, ectopic expression of MYCN in migrating neural crest cells may induce proliferation and maintenance of neural identify yet limit their differentiation potential when they normally reach the ganglia^13^. Some of the key transcription factors that are involved in sympathoadrenal neural crest lineage specification are PHOX2A/B, GATA2/3, HAND2, SOX4/11, and INSM1^15^. Among them PHOX2B, GATA3, HAND2 acts as MTFs in NB in the CRC^5-7,16^.

In this study, we sought to characterize the molecular mechanisms required for reprogramming neuroblastoma cells for differentiation by identifying and validating the master regulators driving self-renewal and those necessary to implement a differentiation program in high-risk neuroblastomas tumors. Understanding disease pathogenesis has the potential to identify clinically relevant targets.

## Results

### All-trans retinoic acid (ATRA) mediated differentiation leads to dynamic changes in super-enhancer landscape of MYCN-amplified NB cells

To identify critical SEs driving transcriptional regulators of NB self-renewal and differentiation, we utilized the well-studied model in which ATRA inhibits NB tumor cell growth and induces differentiation^17,18^. A schematic is detailed in Fig. 1a. Consistent with previous results^17^, ATRA treatment of KCNR cells led to 50-90% decrease in cell growth at 4 and 8 days respectively (Suppl. Fig. S1a). This was accompanied by a 50% decrease in cells in the growth fraction (S+G2/M) of the cell cycle at 8 days (Suppl. Fig. S1b) and morphologic differentiation (Fig. 1b) showing a 4-8 fold increase in neurite length over the time studied (Suppl. Fig. S1c). RNA-seq analysis showed significant downregulation of genes implicated in the NB transcriptional core regulatory complex such as MYCN, ISL1, and GATA3^6^ (Suppl. Fig. S1d, left panel). Gene-set enrichment analysis (GSEA) showed a significant downregulation of Benporath Proliferation and MYCN_UP signatures, and a significant upregulation of Frumm differentiation and Cahoy Neuronal signatures in the transcriptome of cells treated with ATRA (Suppl. Fig. S1d, right panel), confirming ATRA’s ability to influence these pathways in NB.

**Fig. 1:**
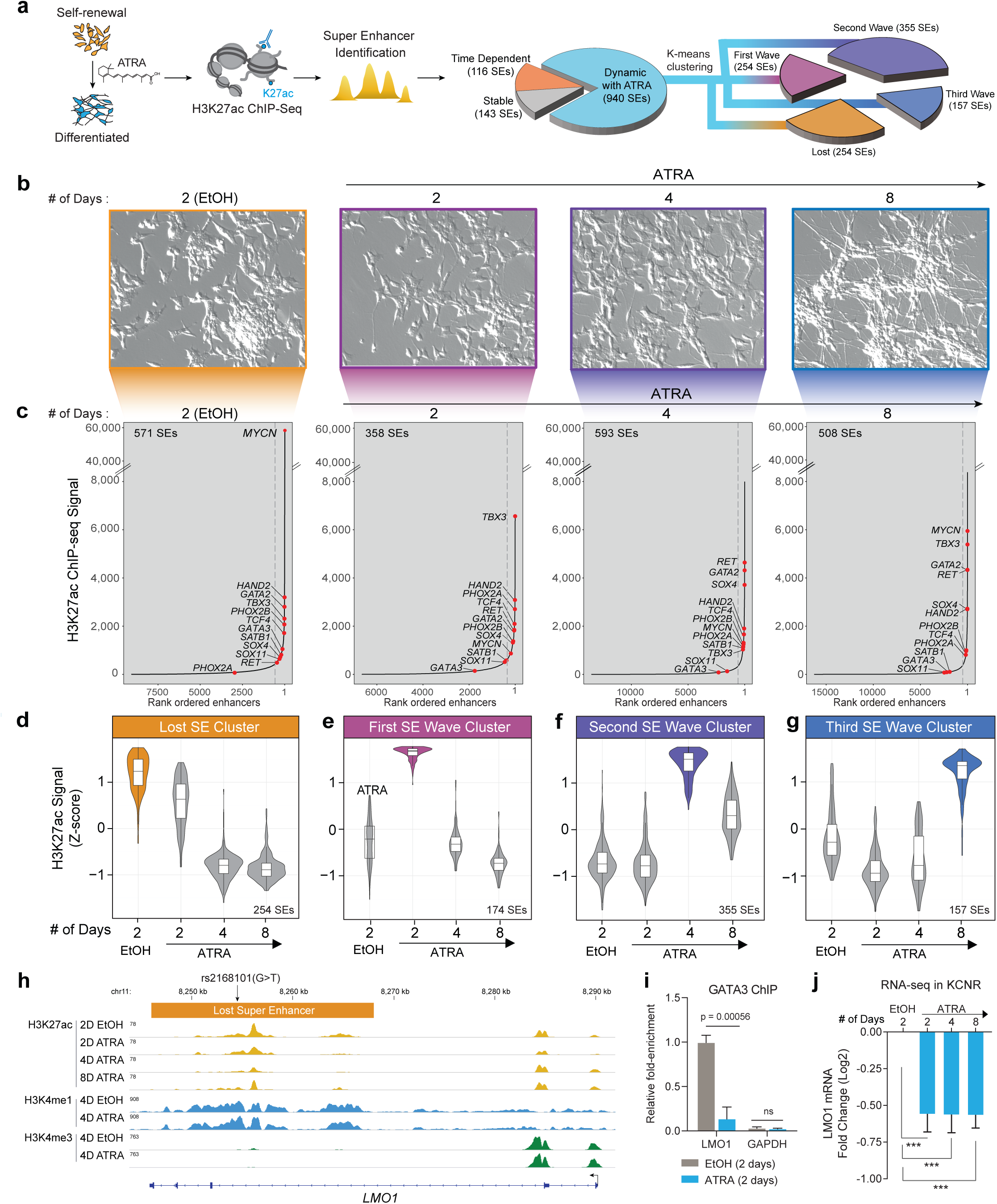
ATRA treatment of NB cells reveals dynamic change in super-enhancer (SE) landscape. a. Schematic representation of the study design and step-by-step approach to identify SEs and downstream clusters after ATRA treatment. SEs were divided into subgroups: Stable with ATRA: SD<30% over 2,4 and 8 days of ATRA treatment; Time Dependent: - Max variance <200 H3K27Ac signal in control (EtOH) and ATRA conditions. The remaining SEs were defined as Dynamic with ATRA and subjected to K-means clustering to identify temporal pattern. 4 clusters of SEs were identified. b. Representative images of NB cells showing time dependent effect of 5μM ATRA treatment on cell morphology. c. H3K27ac binding at super-enhancers ranked by increasing signal. SEs were identified beyond the infection point of increasing H3K27ac load (indicated by grey dashed line). Select SE target genes are highlighted. The SEs were linked to putative target genes by proximity to expressed genes. d-g. Violin plot showing the 4 clusters of SEs identified by z-score normalized H3K27ac signal. The bar on the right shows the number of SEs in that cluster. (d) A group of 254 SEs is highly active in the control or self-renewing cells and is gradually lost in the ATRA treated differentiated cells. (e) A group of 174 SEs is specifically activated 2 Days after ATRA treatment. (f) A group of 355 SEs is specifically activated 4 Days after ATRA treatment. (g) A group of 157 SEs is specifically activated 8 Days after ATRA treatment. h. Representative ChIP-Seq tracks for H3K27ac, H3K4me1 and H3K4me3 in EtOH and ATRA treated cells in KCNR cells, showing loss of LMO1 SE after ATRA treatment harboring the protective rs2168101(G>T) SNP. i. Bar-graph of GATA3 ChIP-qPCR showing relative decrease in GATA3 binding at specific LMO1 SE region following 4 days of ATRA treatment. j. Bar-graph of showing relative decrease in LMO1 mRNA expression.

To gain insight into the changes in the epigenetic landscape due to ATRA mediated differentiation, we performed H3K27ac ChIP-seq at indicated time points on KCNR cells, under self-renewing (EtOH-2D) and differentiating states (ATRA 2D, 4D, and 8D). We defined SEs (Fig. 1c) using a modified ROSE algorithm^19^, to account for the MYCN amplification under all conditions. SEs spanned a median of 11.9 kb (compared with 1.07 kb for typical enhancers) and had 12.7-fold higher load of H3K27ac (median intensity for typical enhancers was 980 compared to 12408 for SEs) (Suppl. Fig. S1e). Total H3K27ac protein levels did not change after ATRA treatment (Suppl. Fig. S1f). The number of typical enhancers identified in the controls (EtOH-2D) essentially remained constant from 2 days (n=9097) to 4 days (n=8872) and decreased 20% by 8 days (n=7239). In the ATRA treated cells there was a dynamic reorganization of the enhancer landscape with the number of enhancers initially decreasing compared to the controls at 2 days but increasing 2-fold from 2 days (n=6661) to 4 days (n=13698) followed by an additional 17% increase by 8 days (n= 16400) (Suppl. Fig. S1g). Similarly, the number of SEs in ATRA-treated cells initially decreased compared to the controls but increased 1.5-fold from 2 days (n=358) to 4 days (n=593) (Suppl. Fig. S1h). The SEs in controls essentially remained constant over time (data not shown).

Consistent with previously published studies, under self-renewal or control conditions (EtOH-2D), we found MYCN acts as the major driver with the most highly active SE followed by the recently identified components of the NB CRC such as *PHOX2B, HAND2, GATA3*^5,19^ (Fig. 1c: EtOH-2D). Upon ATRA treatment, MYCN SE dramatically changed with significant decrease in H3K27ac levels after 2-4 days (Fig. 1c). In addition to MYCN, SE signals associated with multiple genes such as *SOX11, SOX4, GATA3*, and *GATA2* also changed with ATRA treatment (Fig. 1c).

To understand the dynamic changes in the SE landscape mediated by ATRA, we evaluated the temporal patterns of SE changes and identified 1199 SEs (Suppl. Table 1) in KCNR cells under self-renewal and differentiating conditions. All identified SEs were rank-ordered according to their H3K27ac signal. To identify SEs dynamically regulated by ATRA (Fig. 1a), we removed SEs that remained stable over the course of treatment (SD < 30%; n=143) or those whose temporal pattern changed under both control and ATRA conditions (Max variance <200 H3K27Ac; n=116). For the remaining SEs (n=940), K-means clustering based on H3K27ac density z-score was used and identified 4 temporal patterns of regulated SEs (Suppl. Fig. S1i). The temporal pattern of the H3K27ac Z-score for each cluster is plotted in Fig.1d-g. The “Lost” cluster was characterized as a group of SEs (n= 254) whose activity either decreased or was completely lost following ATRA treatment (Fig. 1d). The 3 other SE clusters showed sequential waves of activation in response to ATRA treatment. These were termed 1^st^ SE wave cluster at 2D (n=174), 2^nd^ SE wave cluster at 4D (n=355) and 3^rd^ SE wave cluster at 8D (n=157) (Fig. 1e-g). Using GREAT to infer the biological output of non-coding regions by analyzing annotations of neighboring genes^20^, the Lost SE cluster was found to be enriched in the regulation of signaling pathways and DNA binding followed by stem cell development (Suppl. Fig. S1j). The 1st wave of gained SEs reflected responses to hormones, biomolecules and negative regulation of signal transduction and cell communication while the 2^nd^ and 3^rd^ waves of gained SEs predominantly reflected neural development, cell differentiation and axonogenesis (Suppl. Fig. S1j) consistent with morphological changes and increases in neurite extensions.

Since neuroblastoma arises due to a block in terminal differentiation, we sought to interrogate the dynamically regulated SEs for the presence of neuroblastoma-associated single-nucleotide polymorphisms (SNPs). Genome-wide association studies (GWAS) of neuroblastoma have identified over a dozen susceptibility loci and these genetic associations have implicated genes such as *CASC15, NBAT1, BARD1, LMO1, DUSP12, DDX4, IL31RA, HSD17B12, HACE1, LIN28B, TP53, RSRC1, MLF1, CPZ, MMP20, KIF15*, and *NBPF23* ^21-30^. We intersected NB-associated common SNPs from a recent neuroblastoma GWAS^31^ with the dynamically ATRA-regulated SEs in KCNR cells and observed overlap at the *LMO1* locus on chromosome 11p15 (Suppl. Table 2). *LMO1* is a transcriptional co-regulator that functions as an oncogene in neuroblastoma^16,24^. The most significant SNP, rs2168101 (G>T; p-value 3.18 x 10^−16^, odds ratio 0.70, 95% CI 0.65-0.76) maps to an evolutionarily conserved GATA3 binding site within the LMO1 SE. The risk allele (G) retains the canonical binding site and this SE drives expression of LMO1 ^32^. In contrast, the rs2168101 protective allele (T) ablates GATA3 binding and is associated with low- or no LMO1 expression. The NB cell line KCNR contains the G risk allele associated with tumor formation and harbored a strong LMO1 SE under control (EtOH) condition. Upon ATRA treatment, the LMO1 SE became a Lost SE cluster candidate. Specifically, H3K27ac ChIP-seq indicated a 70% loss of the LMO1 SE after 4 days of ATRA treatment (Fig. 1h). Since the risk allele retains a conserved GATA3 motif, we used ChIP analysis to examine GATA3 binding at this locus and found a 75% decrease in GATA3 binding in this region after treatment with ATRA (Fig. 1i). This was accompanied by a 1.46-fold decrease in *LMO1* mRNA by RNA-seq (Fig. 1j). Taken, together, this suggests a potential involvement of the lost cluster SEs in driving NB tumorigenesis.

### Super-enhancers fluctuate in sequential waves

In each cluster, proximity analysis within topological domains was used to associate SEs with their respective genes (Fig. 1d) and Ingenuity Pathway Analysis (IPA) revealed the predicted cellular localization of the genes. Some 31-37% of SE driven genes in the lost cluster and gained 1^st^ wave cluster encoded proteins that localized to nucleus while in the gained 2^nd^ and 3^rd^ wave clusters, SEs associated with nuclear genes decreased. In these clusters some 40% of the SEs regulate genes whose protein products associate with cytoplasmic processes (Suppl. Fig. S2a). Of the SE associated genes whose proteins localized to the nucleus the greatest fraction encoded transcriptional regulators (Suppl. Fig. S2b).

To identify the transcriptional regulators driving NB self-renewal and those contributing to the differentiated cell phenotype, we focused on the analysis of SEs that associate with transcription factors (TFs) (Suppl. Fig. S2c). SEs driven TFs made up 11% (31/254) of SEs in the lost cluster, while they comprised 9% of the gained 1^st^ wave, 12% of the gained 2^nd^ wave and 9% of the gained 3^rd^ wave of SEs. RNA-seq indicated changes in SEs signal of these TF associated SEs (Suppl. Fig. S2d) corroborated changes in the downstream gene expression (Suppl. Fig. S2e). In the lost cluster the mean expression of TF mRNAs significantly decreased over the course of the 8 days of ATRA (Suppl. Fig. S2e). Similarly, the mean expression of TF mRNAs associated with SEs in the gained 1^st^ and 2^nd^ waves was significantly increased compared to their expression in untreated cells (Suppl. Fig. S2e). Since there were no significant differences in the TFs in the gained 3rd wave cluster over the course of ATRA treatment, this group was not included in further analyses.

To identify the master regulators driving NB self-renewal and those contributing to the differentiated cell phenotype, we focused the analysis on SEs associated with transcription factors (TFs) whose temporal pattern of mRNA expression showed a direct relationship with its linked SE signal, as measured by H3K27ac load at the SE. Since SEs regulate expression of a downstream gene, we focused on genes whose loss or gain of SEs correlated with a decrease or increase in gene expression (r > 0.45) (Fig. 2a, Suppl. Table 3). Decreased SE signal in the lost cluster was associated with a significant decrease in the expression of their associated TFs by 8 days (p < 0.001) (Fig. 2b-c). There were significant increases in expression of TFs associated with the 1^st^ and 2^nd^ gained SEs at 2-4 days (Fig. 2 b,c). Heatmaps showed changes in gene expression of TFs in each cluster (Fig. 2d). The changes in SEs and gene expression were validated in LAN5, another MYCN amplified NB cell line responsive to ATRA. Over 70% of the TFs examined in Fig. 2d showed similar changes in LAN5 at the expression level (Fig. 2e) (Suppl. Fig. S3a, S3b).

**Fig. 2:**
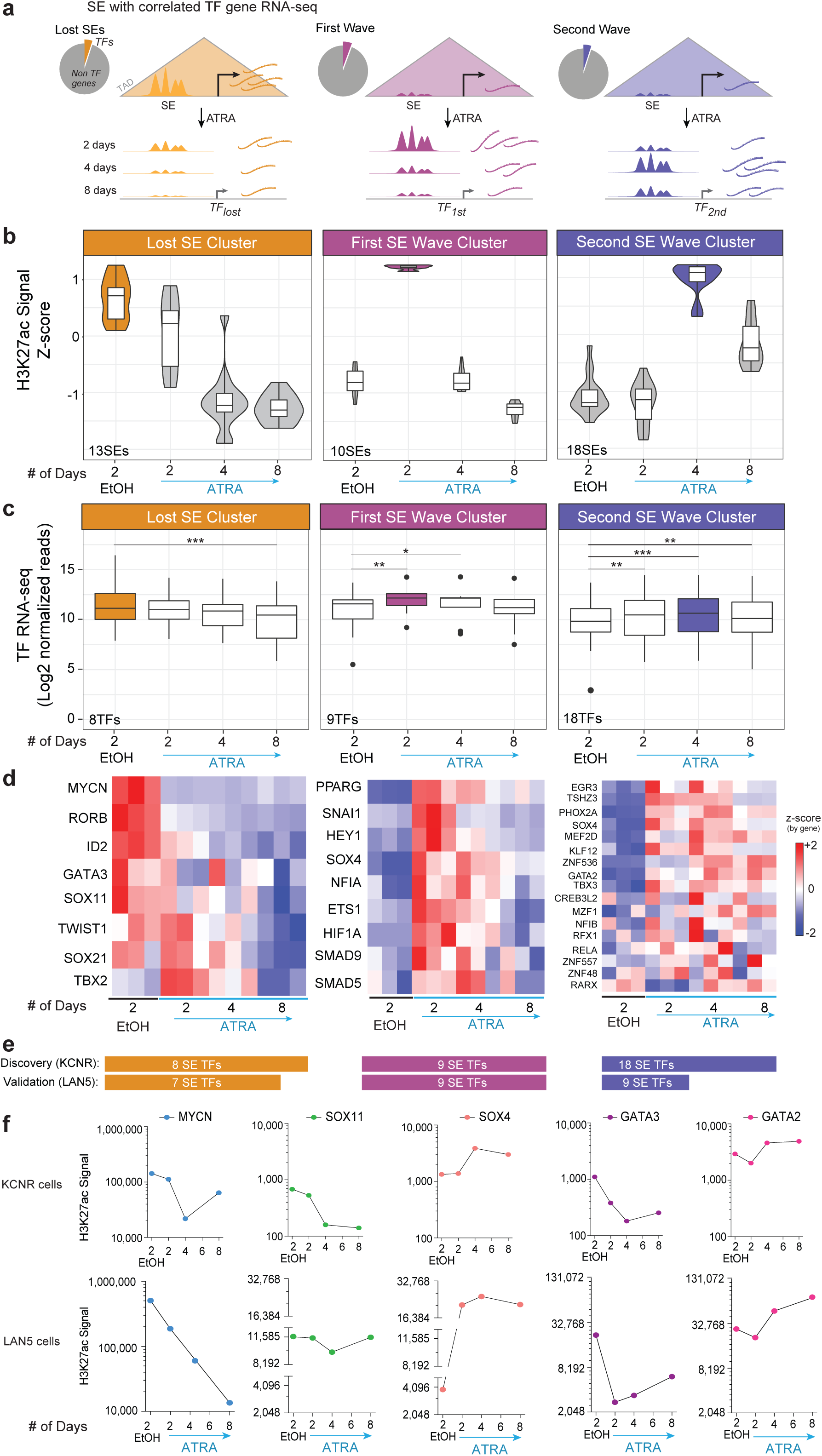
SE driven transcription factors drive proliferation and differentiation of neuroblastoma cells. a. A schematic representation of the analysis steps following SEs identification in each cluster. As TFs drive core regulatory circuitry of a cell, analysis was further restricted to SEs linked only to TFs (Pie charts) and their expression was evaluated at the mRNA level. b. Violin plot showing the 4 clusters of SEs driving TFs (r^2^≥0.45) normalized by z-score of H3K27ac signal. The lost cluster consists of 13SEs, the first wave cluster has 10SEs, and the second wave cluster consists of 18 SEs. c. Bar graph showing expression of the TFs driven by the SEs in each cluster. The 13SEs in the lost cluster drives 8TFs where the loss in SE signal lead to significant decrease in the expression of the corresponding TFs after 8 days of ATRA treatment. Similarly, gain in 10 SEs in the first wave cluster significantly increased expression of its downstream 9 TFs after 2 days and 4 days of ATRA treatment. And 18SEs gained in Second wave led to significant gain in the expression of the downstream 18TFs. d. Heatmap showing expression of individual TFs in each cluster in controls and after 2, 4 and 8 days of ATRA treatment in 3 replicates for each condition. e. Number of SE regulated TFs discovered in KCNR and validated in LAN5. f. Line graph showing dynamic changes at specific SE with ATRA treatment in KCNR and LAN5 cells. H3K27ac signal at the SE associated with MYCN, SOX11 and GATA3 is decreased, whereas signal at SOX4 and GATA2 are increased.

The lost cluster included SEs driving TFs such as MYCN, GATA3, and TBX2 that have been identified as a part of CRC controlling cell growth and proliferation in NB^5,6,19,33^. In addition, this cluster included the SOX11 TF, which is involved in proliferation in sympathetic ganglia at early stages during normal sympathetic nervous system development^34^. We observed a loss in the SE signal associated with MYCN, SOX11 and GATA3 in both KCNR and LAN5 cells (Fig. 2f). Importantly, expression of these TFs also decreased at the protein level in KCNR cells (Suppl. Fig. S2f).

The first and second SE wave clusters included SEs linked to genes such as *SOX4, PPRAG, TBX3, ETS1, HEY1, GATA2, PHOX2A* with increased mRNA expression (Fig. 2d and 2f). Among them, SOX4, GATA2 and TBX3 also showed increase at the protein level (Suppl. Fig. S2f). Fig. 2f shows changes in SEs driving SOX4 and GATA2 in KCNR and LAN5 cells. GATA2 is a potent inhibitor of proliferation of neuronal progenitors and indispensable for the differentiation of several tissues during embryogenesis^35^ and SOX4 expression is increased in late stages of sympathetic nervous system development^34^.

To assess functional effects of the genes in the various clusters, we mined the Project Achilles CRISPR dataset (Cancer Data Science (2018b): Broad Institute Cancer Dependency Map, CRISPR Avana dataset 18Q1 (Avana_public_18Q1) for functional dependencies in the 13 NB cell lines contained within the 391 cancer cell dataset. This analysis showed an enrichment in dependency genes among the 29 TFs driven by SEs in the lost SE cluster (Suppl. Fig. S2g), consistent with a role for these genes in driving self-renewal. Overall, the mean dependency score of the genes driven by dynamically regulated ATRA gained 1^st^, 2^nd^ and 3^rd^ SE waves did not show evidence for dependence (Suppl. Fig. S2g). This suggests that genes with gained SEs after ATRA treatment are not essential for self-renewal or proliferation but rather would be critical to mediate differentiated properties of NB cells.

### SOX family driven regulatory circuits correlate with neuroblastoma prognosis

SEs frequently regulate the expression of MTFs of particular cell lineages and form a CRC, in which the binding motifs for each of the cell type’s MTFs are found in the SEs of that cell type^36^. Because we identified changes in the SE landscape as a result of ATRA treatment, we wanted to investigate if these changes also affected the composition of the CRC. We linked each CRC factor to the SEs activating each other factor, and visualized this using Linked Enhancer Activated Factors (LEAF) plots. Each node (Fig. 3a) represents a transcription factor and each green arrow indicates the presence of a given transcription factor’s DNA binding motif in the active SE regulating the TF to which the arrow points. The size and order of the nodes are proportional to the total mRNA expression of each TF. We found that after ATRA treatment the SOX11 factor is lost from the CRC owing to its lost SE. Interestingly, SOX4 rose to the top of the CRC factors in the presence of ATRA (Fig. 3a). The inward binding of all TFs in the SOX4/11 SEs, inferred from the presence of a given TF’s motif in a SE, decreased for SOX11 and increased for SOX4 upon ATRA treatment (Fig. 3b). The outward binding of SOX4 increased (Fig. 3b) as the abundance of the SOX4 motifs increased in SEs associated with NB cell differentiation (Fig. 3c). In the lost SE cluster, the SE predicted to drive *SOX11* expression had elevated levels of H3K27ac in multiple other NB cell lines, whereas minimal H3K27ac was seen at the *SOX4* SE (Fig. 3d).

**Fig. 3:**
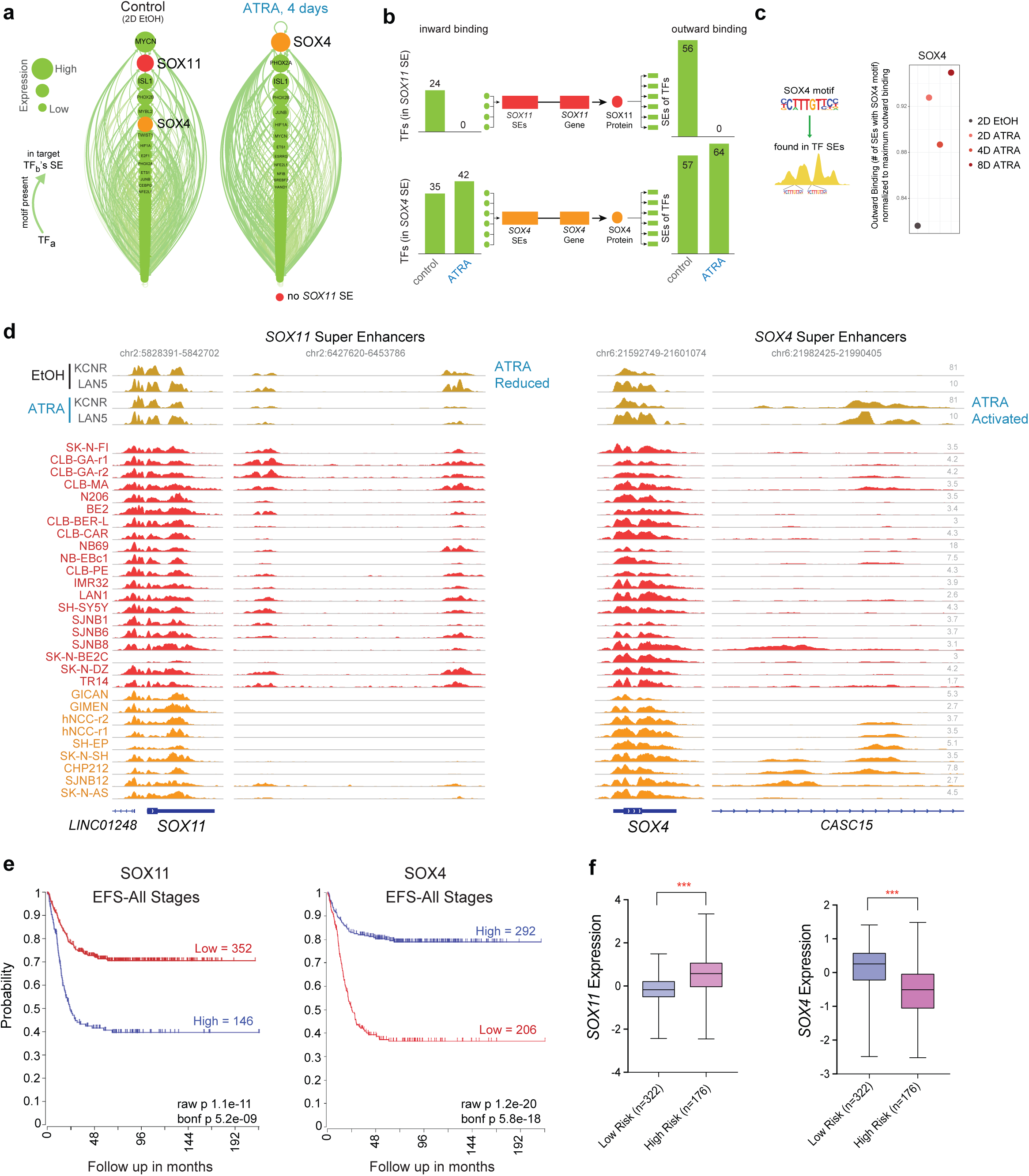
Switching of lineage specific transcription factors induces differentiation in NB cells. a. LEAF (Linked Enhancer Activated Factors) plots of CRC linked by motif presence in their active enhancers, with nodes ranked by mRNA expression in descending order, in controls (EtOH treated) and 4 days of ATRA treatment. b. Inward binding of NB TFs in SOX4 and SOX11 SEs (left), and outward binding in all TF proximal SEs (right), before and after ATRA treatment. Data is from day 4 in KCNR cells, and is similar to connectivity at day 2, and is similar to observed changes in LAN5 cells. c. Dot plot showing increase in SOX4 outward binding (normalized to 1 = maximally outward bound TF) in the SEs in control and ATRA treated samples. d. ChIP-seq track of H3K27ac in ATRA treated KCNR and LAN5 cells (top, gold), compared to group state epigenomics of H3K27ac in other Adrenergic NB cell lines and primary tumors (middle, red) and Mesenchymal NB cell lines and tumors (bottom, orange) at both SOX11 with one of its SEs, and SOX4 with its SE. e. Kaplan-Meier plots based on the expression of SOX11 (left panel) and SOX4 (right panel) in tumors from NB patients at all stages (R2 database: SEQC dataset). Box plot showing significant difference in expression of SOX11 (left panel) and SOX4 (right panel) amongst low and high risk cases. f. RNA expression of *SOX11* and *SOX4* among low and high-risk NB patients. ***P < 0.0001, students t-test with Welch’s correction.

SOX11 and SOX4 are members of the SOX-C family of transcription factors^37^. As sympathoadrenal neural cells differentiate, early expression of Sox11 is followed by increasing expression of Sox4 as Sox11 levels decline^34^. Since little is known of the function of SOX genes in NB, we evaluated the expression of SOX11 and SOX4 in a cohort of 395 normal and tumor tissues sequenced in the TARGET dataset. Relative to the expression in normal tissues we observed higher expression of SOX11 and SOX4 in NB tumor tissues (Sup Fig. 4a). In primary NB tumors (Dataset: SEQC), high levels of SOX11 were associated with a poor outcome (Fig. 3e left panel; Bonferroni p = 1.7e-11), whereas elevated levels of SOX4 were associated with a better prognosis (Fig. 3e right panel Bonferroni p = 5.8e-18). The prognostic significance of SOX11 and SOX4 in primary tumors was replicated in an additional R2 database (Kocak dataset, Suppl. Fig. S4b,c). Assessment of primary NB patients stratified for risk (database: R2 SEQC; n = 498 tumors) reveals increased expression of SOX11 in high-risk cases (Mann Whitney test, p<0.0001), whereas higher expression of SOX4 was found in low-risk cases (Mann Whitney test, p<0.0001) (Fig. 3f).

**Fig. 4:**
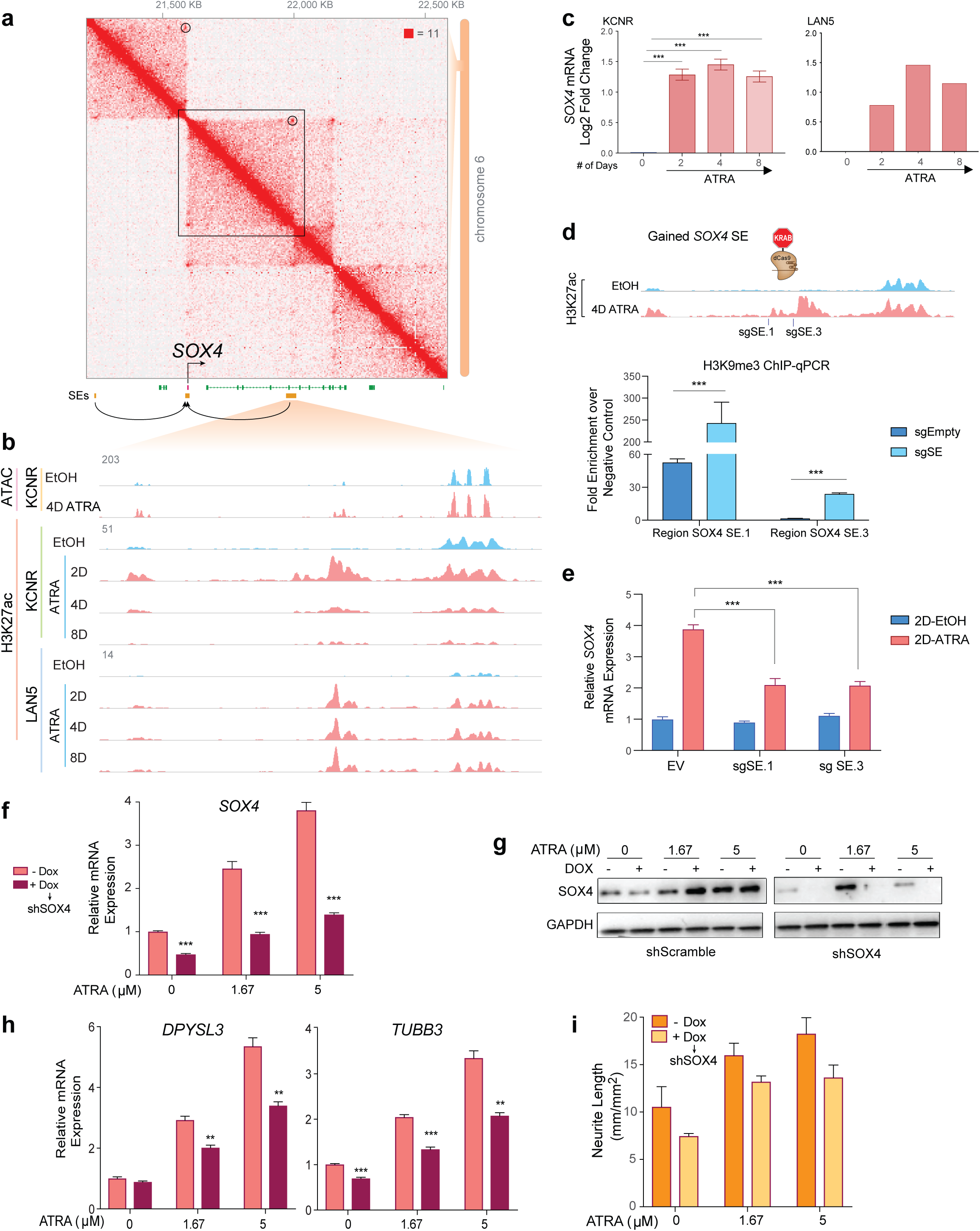
SOX4 silencing leads to inhibition of RA mediated differentiation of KCNR cells. a. Hi-C contact profile surrounding SOX4 locus in KCNR cells with gained SEs shown in both adjacent TADs. b. Zoom in on the SE gained, intronic to CASC15. Representative ATAC-seq and ChIP-Seq tracks for H3K27ac in control (EtOH: blue) and ATRA treated cells (pink), showing increase in H3K27ac peaks at SOX4 SE in KCNR cells. c. Bar-graph showing relative mRNA levels (as normalized reads) of SOX4 after ATRA treatment in KCNR and LAN5 cells. Log2-fold change increase in SOX4 expression was observed post ATRA treatment. In KCNR, data are means ± SD for three replicates; P value generated from paired t test with Welch’s correction. d. Guided suppression of the gained SOX4 SE. Bar-graph of H3K9me3 ChIP-qPCR showing increased H3K9me3 at specific gained SOX4 SE regions following CRISPR-dCas9-KRAB targeting of SOX4 SE. sgRNAs targeting SOX4 SEs were compared to empty vector (sgEmpty), and were performed in KCNR cells. e. Bar-graph showing suppressed SOX4 mRNA induction following CRISPR-dCas9-KRAB targeting of SOX4 SE. sgRNAs targeting SOX4 SEs were introduced into KCNR cells treated with control or ATRA for 2 days. EV: Empty vector. f. qRT-PCR analysis showing relative mRNA levels of SOX4 (left) after doxycycline induced inhibition. 50% inhibition of SOX4 mRNA was obtained after doxycycline treatment. The ATRA mediated increase in SOX4 levels were also reduced ∼60-70% on doxycycline treatment. g. Western blots showing DOX induced downregulation of SOX4 (via shRNA SOX4, right) during ATRA induced expression of SOX4 (with shRNA scramble control on the left). h. SOX4 reduction slowed ATRA induced activation of differentiation marker genes *DPYSL3* (left panel) and *TUBB3* (right panel). i. Bar-graph showing differentiation index as measured by neurite length. SOX4 silencing reduced ATRA mediated differentiation of KCNR cells.

### Gained cluster super-enhancer contributes to increased *SOX4* mRNA

Having observed the gain of the *SOX4* SE and increases in *SOX4* expression with ATRA, we performed Hi-C in KCNR cells to evaluate the prediction of *cis-*interactions between the SEs and the *SOX4* promoter and showed that they occur within the same TAD boundaries (Fig. 4a). We evaluated changes in the chromatin accessibility in this region by ATAC-seq and did observe chromatin opening in the newly gained SE region after ATRA treatment (Fig. 4b). The SE linked to *SOX4* gained H3K27ac signal after ATRA treatment (Fig. 4b) was associated with a 3-fold increase in mRNA (Fig. 4c-left panel). Similar observations were made in the LAN5 NB cell line where ATRA treatment led to a gain of the *SOX4* SE (Fig. 4b) that was associated with a 3-fold increase in mRNA (Fig. 4c-right panel).

To assess the functional contribution of this SE to *SOX4* mRNA expression, we targeted the SE with 2 sgRNAs, using the dCas9-KRAB one-vector system. The KRAB domain leads to reversible silencing of the chromatin by deposition of H3K9me3 modifications at the loci targeted by the dCas9^38^. We evaluated changes in *SOX4* expression after 2 days of ATRA treatment (Fig. 4d, top panel) and observed that the sgRNAs led to 2.5 to 13-fold increase in H3K9me3 signal over the negative control at the specific sgRNA loci compared to minimal or no enrichment for the empty vector (Fig. 4d, bottom panel). This was accompanied by significant decrease (p<0.001) in the ATRA-mediated upregulation of *SOX4* expression (Fig. 4e) demonstrating that the physically linked *SOX4* SE regulates *SOX4* mRNA expression. No changes of *SOX4* mRNA levels were observed in the control cells between negative sgRNA and SE sgRNA transduced cells (Fig. 4e), which could be due to the extremely low basal level of H3K27ac signal at this region.

As SOX4 increases with differentiation of NB cells, we tested whether the loss of SOX4 expression would affect NB differentiation associated genes. To do this we generated a doxycycline inducible SOX4 knock-down cell line. Doxycycline treatment led to at least a 50% decrease in *SOX4* mRNA (Fig. 4f) and protein levels (Fig. 4g). Inhibition of SOX4 expression attenuated the ability of ATRA to induce differentiation markers DPYSL3 (Fig. 4h, left panel) and TUBB3 (Fig. 4h, right panel) and resulted in a 20% decrease in ATRA induced neurite extension (Fig. 4i). These data indicate SOX4 acts to enhance ATRA mediated differentiation in neuroblastoma.

### Lost cluster super-enhancer drives SOX11 and contributes to NB self-renewal

As we observed the gain of SOX4 coincided with the loss of SOX11 from its dominant position in the CRC during ATRA induced differentiation, we evaluated if the SEs distal to *SOX11* were indeed driving it directly. The SOX11 linked SE was lost after ATRA treatment (Suppl. Fig. S5a) with an 80% decrease in H3K27ac signal and was accompanied by significant decrease in SOX11 mRNA (p < 0.001) (Suppl. Fig. S5b) and protein levels (Suppl. Fig. S2f). To evaluate *cis-*interactions between the SEs and the *SOX11* promoter, we performed Hi-C in KCNR cells and observed *SOX11* and its SEs are encircled within the same TAD boundaries (Fig. 5a). This provided direct evidence linking the bioinformatically-associated *SOX11* SE to the *SOX11* TSS. To assess the functional contribution of this SE to *SOX11* mRNA expression, we targeted the SE with three sgRNAs (Fig. 5b, upper panel) using the dCas9-KRAB system. In 2 of 3 sgRNAs, we observed a 50-80-fold enrichment in the H3K9me3 signal over negative control at specific sgRNA loci, compared to no enrichment for the empty vector at the targeted *SOX11* SE (Fig. 5b, lower panel). This was accompanied by a 1.5-2-fold decrease in H3K27ac at the same locations (Fig. 5c). Targeted chromatin modification swapping at the *SOX11* SE resulted in a significant decrease in *SOX11* expression (p < 0.001) (Fig. 5d). These results functionally confirmed this lost SE directly regulates *SOX11*.

**Fig. 5:**
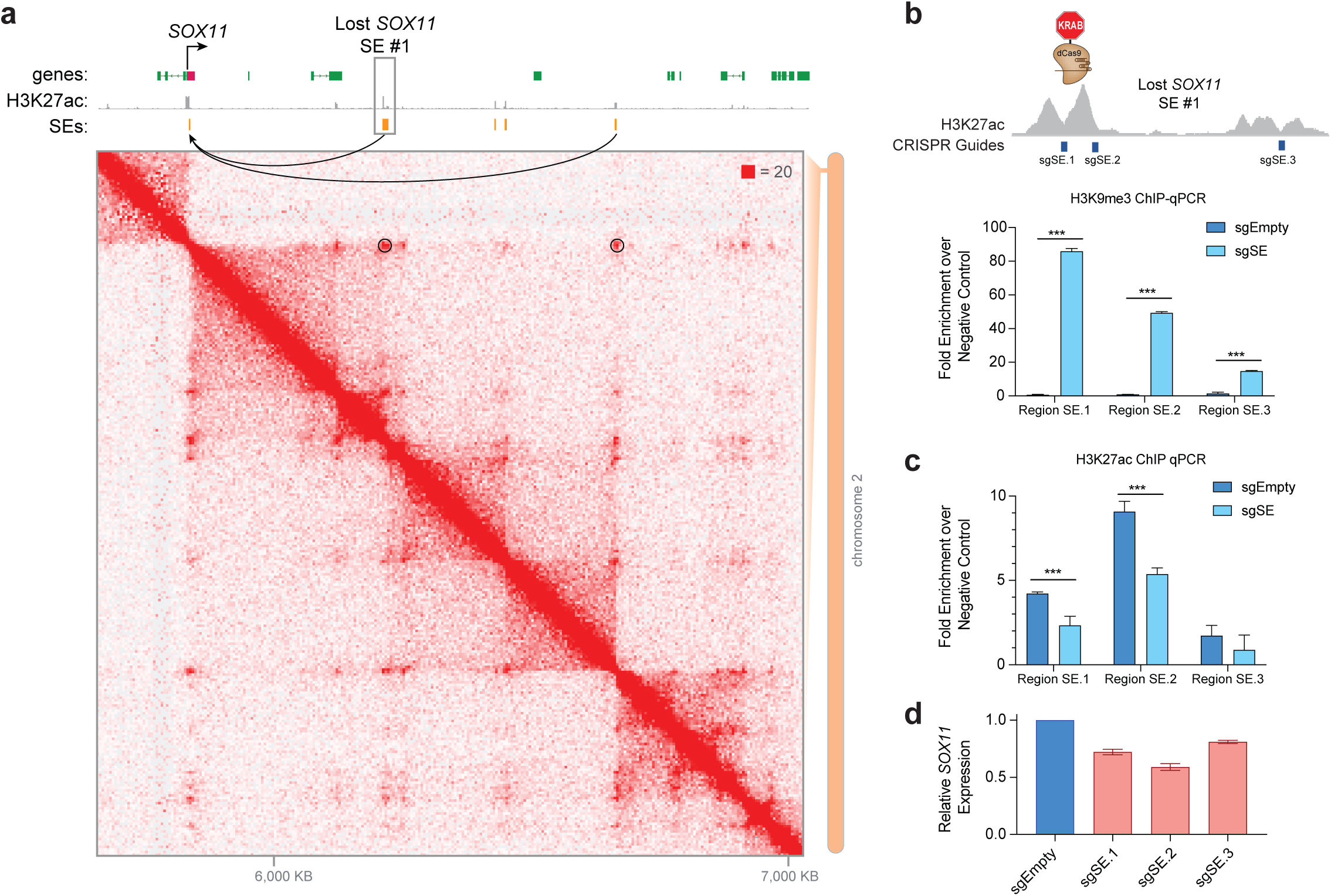
SOX11 expression is regulated by SEs in NB cells. a. Hi-C contact-map surrounding SOX11 locus in KCNR cells. Two lost SEs connected to the SOX11 TSS are highlighted, with SOX11 gene and the largest lost SE (#1) labeled. b. Upper panel: Guided suppression of the lost SOX11 SE. Lower panel: Bar-graph of H3K9me3 ChIP-qPCR showing relative increase in H3K9me3 peaks at specific SOX11 SE regions following CRISPR-dCas9-KRAB targeting, compared to empty vector (sgEmpty) in KCNR cells. c. Bar-graph of H3K27ac ChIP-qPCR showing relative decrease in H3K27ac at specific SOX11 SE region following CRISPR-dCas9-KRAB targeting. sgRNAs targeting SOX4 SEs led to significant decrease in H3K27ac peaks compared to empty vector (EV) in KCNR cells. d. SOX11 mRNA levels (measured by RT-qPCR) decreased upon CRISPR-dCas9-KRAB mediated repression of SOX11 SEs in KCNR cells.

### SOX11 inhibition decreases cell growth in NB cells

Since SOX11 is highly expressed in NB cells and tumors, we evaluated whether NB is dependent on SOX11 for self-renewal. We first analyzed the genome wide CRISPR library screen of 391 cancer cell lines^6^ and found that among various tumor types, NB cells are most selectively dependent on SOX11 (Fig. 6a), although there are a range of dependencies among different NB cell lines (Suppl. Fig. 5c) in which MYCN amplification appears to predispose cells to SOX11 dependence. The siRNA inhibition of SOX11 caused a 90-100% inhibition of SOX11 mRNA and protein levels (Fig. 6b) and a significant decrease in relative cell number in KCNR cells (Fig. 6c). Inhibition of SOX11 also resulted in significant decreases in the relative growth as measured by confluence, in 4 of 6 additional NB cell lines tested (Fig. 6d).

**Fig. 6:**
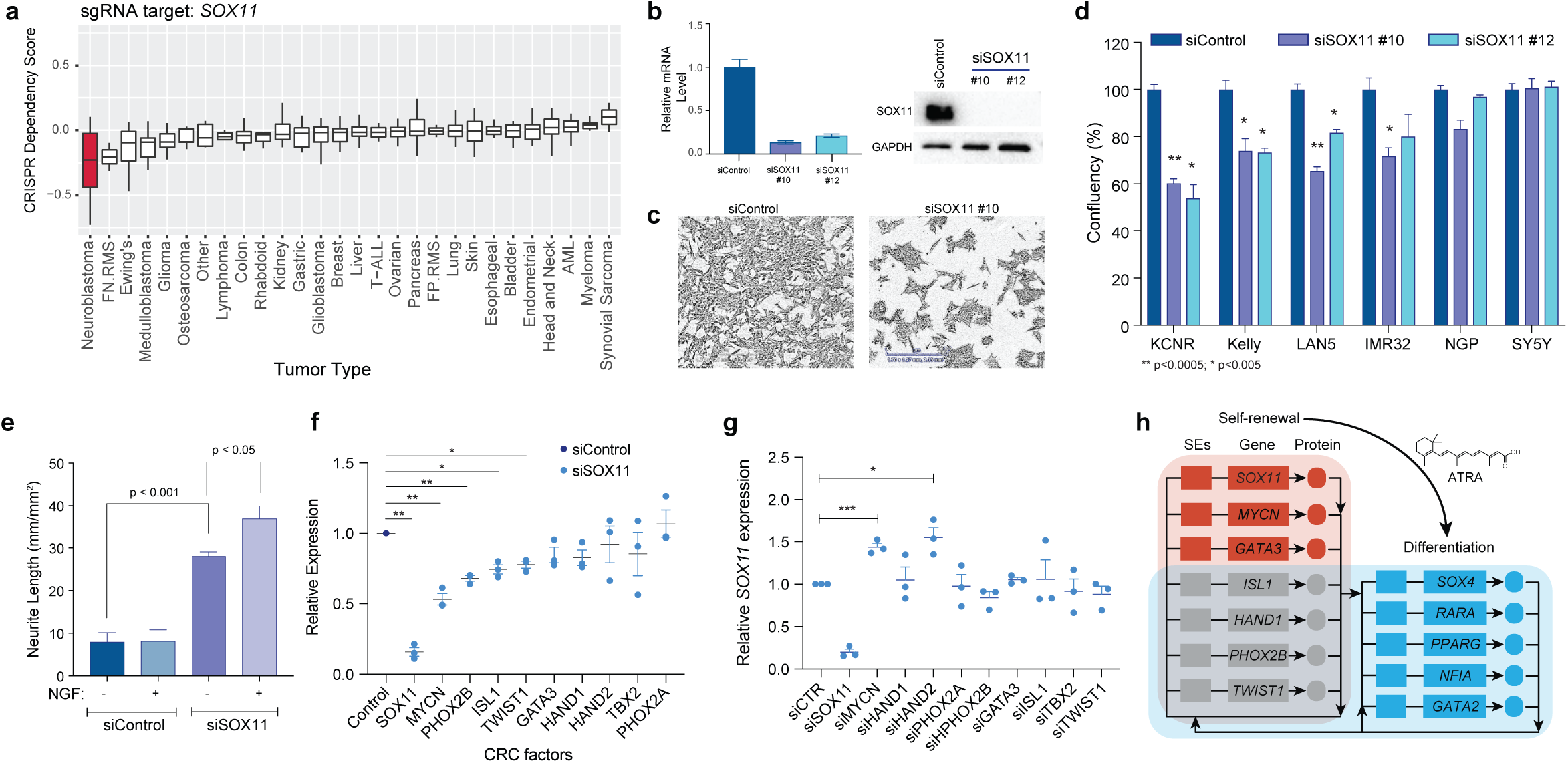
SOX11 drives cell growth and proliferation of neuroblastoma cells. a. Bar graph showing dependency of different cancer cells -on SOX11 for cell growth and proliferation. NB cells are most preferentially dependent on SOX11 for growth and proliferation compared to other tumor types. Data mined from Project Achilles genome wide CRISPR-Cas9 screen. b. qRT-PCR analysis showing relative mRNA levels of the SOX11 after silencing for 48 hr. Bars show the mean ± SEM of three replicates. The adjacent western blot shows inhibition of SOX11 at the protein level. c. Representative images of KCNR cells following genetic silencing of SOX11. d. Multiple NB cells demonstrate significant decreased cell growth after silencing of SOX11. Graph showing representative data from at least 3 biological replicates. e. Significant increase in differentiation index as analyzed by neurite length was observed after SOX11 inhibition, which was further increased upon NGF treatment. f. RT-qPCR analysis showing relative mRNA levels of known NB regulatory TFs after silencing of SOX11 for 48hr. Graph showing representative data from 3 biological replicates. g. RT-qPCR showing relatively small *SOX11* mRNA changes upon knockdown of other CRC members. Each TF was knocked down for 72 hrs. in KCNR cells, showing representative data of 3 biological replicates. h. Model of CRC network swapping during ATRA induced differentiation of NB. Genes highlighted were those which were consistently called in KCNR and LAN5 cells, both by enhancer analysis and gene expression changes. Lines represent inferred or verified binding by ChIP-seq, and do not necessarily indicate that such SE binding is an essential positive regulation event.

To gain further insight into the potential mechanisms contributing to SOX11 mediated growth inhibition and differentiation in NB cells, we evaluated genome-wide changes in the transcriptome by RNA-seq (Suppl. Fig. 5d). We found that the genes regulated by SOX11 were involved in nervous system growth and development (Suppl. Fig. 5e, upper panel) and cellular growth and proliferation (p value 8.05E-06 – 1.21E-23) (Suppl. Fig. 5e, lower panel) amongst other molecular functions (Suppl. Table 4). Importantly, knockdown of SOX11 significantly increased NTRK1 mRNA levels (Suppl. Fig. 5f) and elevated levels of NTRK1 are found in the tumors of NB patients that have a good prognosis^39,40^. Inhibition of SOX11 caused an increase in differentiation-associated phenotype of neurite extension (Fig. 6e), an effect that was further amplified by NGF.

Cancer cells have a remarkable dependency on the transcription factors that form an autoregulatory CRC, where each member directly regulates the expression of its own gene as well as those encoding the other CRC transcription factors^5-7^. To determine the impact of SOX11 on other components of the CRC, we evaluated the expression of the other CRC TFs after SOX11 knockdown. Indeed, decreased SOX11 expression led to decrease in CRC targets like MYCN, PHOX2B, ISL1 and TWIST1 in KCNR cells (Fig. 6f). However, silencing of NB CRC components GATA3, PHOX2B, ISL2 and TBX2 (Suppl. Fig. 5g) did not lead to decreases in SOX11 (Fig. 6g) suggesting that SOX11 may be so well supported by TF binding at its SE that loss of any one of these CRCs individually was not detrimental to its expression. Loss of multiple CRC members at multiple SEs is likely needed to stop SE activation, keeping in mind that dCas9-KRAB inactivation at one SE constituent bound by NB CRCs downregulated *SOX11* (Fig. 6d). That silencing of MYCN leads to increases in SOX11 suggests that SOX11 functions upstream of MYCN which exerts feedback inhibition on SOX11 expression. This idea agrees with recent evidence that CRC interactions are not always feed-forward for each TF and each SE^41^. Our data taken together, suggests that SOX11 sits atop the CRC hierarchy in self-renewing NB, and its displacement in some NB cell lines is important for rewiring to a differentiated state. This model (Fig. 6h) illuminates the cell-identity determining circuits responsible for giving retinoic acid such potent antitumor and pro-differentiation capacity in the treatment of NB patients.

### Discussion

In this study we identify specific temporal and dynamic spatial regulatory events involving SE driven genes that propel neuroblastoma cells from a self-renewal/proliferative state to a differentiated state. We find there is an initial loss of a cluster of SEs driving processes associated with stem cell development, DNA binding and kinase activity followed by the sequential gain of processes regulating signaling events that ultimately lead to the gain of SEs driving processes regulating neural differentiation. By focusing on SEs driving TFs we find the cluster of lost SEs drive many of the sympathoadrenal lineage specifying TFs integral to the NB CRC^5,16,33,42^. However, we also identify SEs regulating *SOX11* and *SOX4* expression that are coordinately lost and gained during the differentiation process. The loss of the SE driving *SOX11* expression decreases NB cell growth making NB cells more responsive to differentiation signals. The gain of SE increases SOX4 expression and is associated with the enhanced differentiation state as functional studies showed decreases in *SOX4* expression attenuates ATRA-mediated differentiation.

Our study uncovers an important role for SOX11 and SOX4 in NB biology. SOX genes encode TFs that bind DNA through conserved high-mobility group domains and have been shown to function as pioneer TF that have important roles in developmental processes regulating cell fate and differentiation. In neural cells the orchestrated yet transient expression and activity of SOX11 is crucial for the precise execution of a neurogenic program^43,44^, it is needed to maintain progenitor pools and promote neuronal differentiation^45,46^. The sequential expression of different SOX genes has been shown to be important for early specification of the neural lineage in ES cells with the subsequent expression of different SOX genes required for the transition to later stages of neural differentiation^43^. In developing sympathetic ganglia, Sox11 is essential for neuronal proliferation but during development as SOX11 expression decreases, levels of SOX4 increase^34^. Little is known about the role of SOX genes in NB biology. In this study, we find sequential changes in SEs driving SOX gene expression upon NB cell differentiation. Initially SOX11 is expressed and contributes to their maintenance in a self-renewal or proliferating progenitor state. Upon differentiation, the SE driving SOX11 expression is lost while SE driving SOX4 levels is gained. Silencing of SOX11 is associated with an increase in the expression of differentiation genes making NB permissive for differentiation but requiring additional stimuli such as the ATRA-induced gain of the SE driving SOX4 (Fig. 2), environmental factors such as NGF (Fig. 5) or other genes (Fig. 2). Our ability to document the sequential nature of the SOX11 and SOX4 interactions at a genome wide level has been limited by an inability to reliably ChIP either SOX11 or SOX4 with currently available reagents. However, the loss of SOX11 binding motifs and subsequent gain of SOX4 binding motifs in SEs associated with NB cell differentiation (Fig. 3a-c) supports such the sequential involvement of SOX genes model of differentiation.

While this study has focused on the functional consequences of SOX11 and SOX4 during NB differentiation, we believe other genes such as GATA3 and GATA2 also play key roles during this differentiation process. The SE driving GATA3 is also lost while the SE driving GATA2 is gained during NB differentiation. GATA2 is a potent inhibitor of proliferation of neuronal progenitors and indispensable for the differentiation of several tissues during embryogenesis^35^. Studies in other cancers show that depending on the type of cancer and cellular context, SOX4 functions as a tumor suppressor, exerting its effects through the induction of apoptosis^47,48^. Higher levels of GATA2 and SOX4 are found as NB cells differentiate and in the tumors of NB patients’ high levels of GATA2 or SOX4 are significantly associated with a better prognosis.

In the companion report, Zimmerman *et al.* independently observed that retinoic acid causes both loss of adrenergic CRC members (such as MYCN, PHOX2B, and GATA3), and gained new CRC members (RARA, SOX4 and TBX3). While our study focuses on functional roles of key CRC members SOX11 and SOX4, the related work explores another important question: why does ATRA fail to differentiate a subset of MYCN-or MYC-driven NB? One scenario where ATRA does not suppress MYCN or MYC, and thus cannot reprogram the adrenergic, arises from HAND2 super-enhancers being hijacked to control the expression of MYCN or MYC. This is in line with the reproducible finding across both studies that *HAND2* super-enhancers are not perturbed by ATRA, as HAND2 is a member of both the self-renewal and differentiated CRCs. Thus, in the case where NB patient tumors harbor translocations between HAND2 SEs and the MYCN or MYC oncogenes, retinoid-based treatments would not be effective in triggering differentiation and prevention of tumor relapse. This could potentially inform clinical decisions for therapy.

The cluster of lost SEs are linked to many genes involved in NB pathogenesis. LMO1 is known to synergize with MYCN to generate NB in zebrafish^49^ and also functions as a key coregulator of the neuroblastoma CRC^16^. The SE driving GATA3 expression is in cluster lost upon differentiation. The consequences of this GATA3 decrease impacts LMO1 as the NB risk allele SNPrs2168101 G>T which creates a novel GATA3 binding site leading to a SE driving the transcriptional regulator LMO1 (ref ^32^) is lost leading to decreased LMO1 expression. The lost cluster also contains SEs linked to 8 TFs whose expression are significantly downregulated upon NB cell growth arrest and differentiation. The involvement of these TFs is supported by the findings that 1) high expression of 6 out of the 8 genes (MYCN, ID2, TBX2, TWIST1, SOX21, and SOX11) is associated with poor prognoses in primary NB tumors (R2 database; SEQC dataset); 2) several are key components of or linked to the NB CRC (GATA3, TBX2, MYCN)^5,6,33^ and 3) many function as NB dependencies or are known to regulate NB growth and differentiation. TWIST1 is a direct target of MYCN^50^ and regulates expression of the MYCN enhancer axis^51^ while ID2, also a known inhibitor of neural differentiation in neuroblastoma and is regulated by MYCN^52,53^. Our results indicate that these genes contribute to the self-renewal capacity of NB cells as the loss of their SE reduces their expression and is critical for cells to fully implement a post-mitotic differentiation program.

ATRA treatment leads to decrease in SOX11 expression by the loss of the SE driving SOX11. We find that SOX11 serves as an input to the NB Core Regulatory Circuit (CRC) stimulating expression of MYCN and CRC members but is not itself dependent on any other individual TF of the CRC but may be dependent on multiple CRC members. Our finding that SOX11 stimulates MYCN but MYCN inhibits SOX11 expression suggests that SOX11 functions upstream of MYCN in a transcriptional regulatory path in which induction of a downstream target leads to feedback suppression of the upstream regulator. Thus, SOX11 functions via MYCN as an input into the NB CRC (Fig. 6h). As the NB cells are rewired for differentiation by ATRA we find 5 gained SEs driving differentiation associated genes that are common between the 2 NB cell lines examined (Fig. 6h). While we have studied a functional role for SOX4 in this study, we do not believe SOX4 functions alone. The role of GATA2 and other genes will be explored in future studies. The recent finding that SOX11 may function as a pioneer TF^54^ raises the possibility that SOX11 may have a unique function in NB cells developing along a sympathoadrenal differentiation pathway.

In summary, we have identified distinct groups of cell-state specific SEs driving processes consistent with changes needed to switch a self-renewal program to a differentiation program. In the future, it will be exciting to explore wider implications of SE circuitry creating potential energy barriers (Waddington mountains) locking cancer cells in de-differentiated states. The influence of small molecules on reshaping this landscape (as here) remains largely unexplored but will be a fruitful space to map the mechanisms by which cell identity is influenced by chemical and extracellular signals.

## Acknowledgements

This work was funded by the Center for Cancer Research, Intramural Research Program at the National Cancer Institute, NIH. We would like to thank Drs. Sridhar Hannenhalli, and Marielle Yohe for review of the manuscript. We are grateful to Drs. Jack Shern, Laura Kerosuo and Mark Zimmerman for helpful discussions. We also thank Drs. Xinyu Wen, Haiyan Lei and Xiyuan Zhang for advice on bioinformatic analyses. We thank Bao Tran, Jyoti Shetty, and Yongmei Zhao from NCI Sequencing Facility for ChIP-DNA and RNA sequencing.

## Materials and Method

### Cell Lines

Human MYCN amplified neuroblastoma (NB) cell lines [SMS-KCNR (KCNR), LAN5, IMR32, Kelly, and NGP] and MYCN-WT NB cell lines [SH-SY5Y (SY5Y)] were used in this study. NB cell lines were obtained from the cell line bank of the Pediatric Oncology Branch of National Cancer Institute and have been genetically verified. Cells were cultured in RPMI-1640 medium supplemented with 10% fetal bovine serum (FBS), 2mM L-glutamine, 100 μg/mL streptomycin and 100U/ml penicillin at 37°C in 5%CO_2_. Human embryonic kidney cells (HEK293T) were obtained from ATCC and cultured in Dulbecco’s modified Eagle’s media (DMEM) with 10% FBS, 100 μg/mL streptomycin and 100U/ml penicillin. For control or SOX4 stable knockdown, SMARTvector Inducible Lentiviral shRNA (GE Healthcare) particles were generated using HEK293T cells. Stable knockdown KCNR cell line was generated by infecting KCNR cells with either non-targeting control shRNA lentiviral particles (SMARTvector Inducible Lentiviral shRNA GE Healthcare, Cat # VSC11707) or SOX4 shRNA lentiviral particles (Cat # V3SH11252-226882070, 227115611, 227170688) and selected using puromycin (500ug/ml, Sigma-Aldrich) to generate pools. Doxycycline (Dox, 0.125ug/ml) was used to induce shRNA expression. For stable CRISPR mediated super-enhancer knockdown KCNR cell lines, the sgRNAs (Suppl. Table 5) were cloned in pLVhU6-sgRNA hUbC-dCas9-KRAB-T2a-Puro (Addgene # 71236) ^38^ using Esp3I cut site and sequenced for positive clones. Cells were transduced and selected as described above. Target inhibition was validated using RT-PCR 7 days after infection. All cell lines were routinely tested for and were free of mycoplasma.

### Differentiation Assay

All-*trans* retinoic acid (ATRA) (Sigma Aldrich (R2625)) dissolved in 95% ethanol was used as a differentiating agent. KNCR and LAN5 cells were plated at a density of 3.5×10^6^ cells/150mm dish and treated with ethanol or ATRA (5μM). After 2, 4 and 8 of days treatment, media was decanted and cells washed twice with ice-cold PBS and detached mechanically from the culture dish. Viable cell number was determined by trypan-blue exclusion and collected as separate aliquots for protein, RNA and FACS analysis as previously described ^55^. For ChIP, cells were fixed at indicated time points, with 1% Formaldehyde solution (Thermo Scientific, Waltham, MA). For ATAC-seq, an aliquot of 50,000 cells under each condition was made. Cell confluency and neurite extension assays using Essen IncuCyte ZOOM or FLR were adopted to evaluate cell proliferation and differentiation in real-time.

### Transient Transfection

For transient inhibition of transcription factors, NB cells were electroporated, using Nucleofector Solution V, Program A-30 for KCNR, LAN5, Kelly, NGP and SY5Y cells and Nucleofector Solution L, Program C-005, in a Nucleofector II device (Amaxa Biosystems), with ON-TARGETplus siRNAs (Suppl. Table 6), according to the manufacturer’s instructions. On-target plus non-targeting siRNA (5’-UGGUUUACAUGUCGACUAA-3’, Dharmacon) was used as a control. The SMARTpool and 4 independent SOX11 siRNA were tested (Suppl. Table 7), out of which 90% inhibition was achieved for 2 siRNA (#10 and #12) and were used for downstream studies.

### RNA isolation and RT-PCR

For RNA extraction approximately 2×10^6^ cells were collected per biological replicate. Total RNA was isolated from NB cells after ethanol or ATRA treatment; siRNA inhibition, SOX4 knockdown and SE knockdown, using RNeasy Plus Mini kit (Qiagen Inc.), as per the manufacturer’s protocol and analyzed for integrity using an Agilent 2200 TapeStation System (Agilent Technologies). Independent replicates (2-3) were performed for each group.

For quantitative PCR analyses, preparation of cDNA (ABI, Cat # 4387406) and real-time qPCR using Fast-SYBR green mix (ABI, Cat # 4385612) was done according to manufacturer’s protocol. Quantitative measurements of genes’ levels were obtained using the BIO-RAD CFX Touch Real-time (RT) PCR detection system and performed in triplicate. Ct values were normalized to HPRT levels. Representative data from biological replicates were shown in this study. Gene expression as quantified by RT-PCR is expressed as relative expression compared to the basal expression under control or self-renewing state. The primers used for qPCR are listed in Suppl. Table 8. In doxycycline induced shSOX4 cell lines, *SOX4* levels were inhibited 50% after 4 days of DOX treatment. Following this, cells were treated for 4 days either with ethanol or ATRA and gene expression was analyzed.

### RNA-sequencing

#### Sample preparation and sequencing

Total RNA was isolated and subjected to RNA-seq analysis from KCNR and LAN5 cells after 2, 4 and 8 days of ethanol or ATRA treatment and KCNR cells transfected with siControl and siSOX11 (#10) for 48hhrs. Total RNA was extracted using RNeasy Plus Mini kit (Qiagen Inc.), according to manufacturer’s protocol. Strand-specific whole transcriptome sequencing libraries were prepared using TruSeq® Stranded Total RNA LT Library Prep Kit (Illumina, San Diego, CA, USA), following the manufacturer’s procedure. RNA-seq libraries were sequenced on a HiSeq2500for paired-end with read length of 126 bp, or HiSeq 3000/4000 for paired-end with read length of 150 bp.

#### Data Processing and analysis

Briefly, the Fastq files were processed using Trimmomatic (version 0.30) ^56^ to remove low quality bases. The trimmed fastq data were aligned to human genome hg19 with STAR (version 2.4.2a) ^42^ and annotated using UCSC or Ensemble file version 19 (Ensembl 74) using Partek® Flow® software, version 7.0 Copyright ©; 2020 Partek Inc., St. Louis, MO, USA. STAR software also generated the strand-specific gene read counts. About 77% of 70 million reads per sample were mapped to human genome uniquely for a total of 90% mapping rate. To eliminate batch effect and identify differential gene expression, the gene reads count data from STAR were analyzed with R Package Deseq2 ^57^. Deseq2 uses methods to test for differential expression by use of negative binomial generalized linear model and was also used to normalize the reads count data to generate z-scores for heatmap display. Heatmaps were created using heatmap.2 function in g plots (version 2.17.0).

#### Pathway analysis

Statistical results of differentially expressed genes from Deseq2 were analyzed using QIAGEN’s Ingenuity® Pathway Analysis (IPA QIAGEN Inc.) Genes with false discovery rates (FDRs) <0.05 were used as input for IPA core analysis that identified significant canonical pathways.

#### Gene Set Enrichment Analysis

Enrichment for curated gene sets (Molecular Signatures Database v 7.0, http://software.broadinstitute.org/gsea/msigdb/) was performed using GSEA software version 4.0 ^58,59^. Differential expression output from Deseq2 (comparing RNA-seq from perturbation with ethanol verses ATRA, or control siRNA verses SOX11 siRNA) was used to rank genes according to the log2 fold change. The GseaPreranked tool was employed with 1000 permutations, and only gene sets with a maximum list size of 500 were considered.

#### Protein isolation and western blot analysis

Protein lysates from with or without ATRA treated cells; siControl and siRNA inhibited cells, shControl and shSOX4 cells were extracted using RIPA buffer (50mM Tris pH 8.0, 150nM NaCl, 0.1% SDS, 0.5% Sodium deoxycholate, and 1% Triton-X 100), containing 1X Protease and Phosphatase Inhibitor cocktail (Thermo Fisher, 78442). Protein concentration was measured using Bradford Assay (Biorad, Hercules, CA, 5000006). Protein (15-30μg) was denatured in 4X Laemmli Sample Buffer (Biorad, Hercules, CA, #1610747), separated on 4-20% precast SDS gel (Biorad, Hercules, CA), and transferred on a nitrocellulose membrane. Membranes were incubated in a blocking buffer [5% dry milk in TBST (TBS with 0.2% Tween-20)] for 1 hr followed by overnight incubation with the primary antibody in blocking buffer at 4°C. Bands were visualized using enhanced chemiluminescence (Amersham Biosciences) or SuperSignal™ substrate (ThemoFisher). The antibodies and the dilutions used in the study are listed in Suppl. Table 9.

#### Cell Cycle analysis by FACS

For determining changes in cell cycle progression before and after ATRA treatment, 2-5 million NB cells were harvested and fixed by resuspending them in an ice-cold solution containing 0.5 mL phosphate-buffered saline (PBS) and 4.5 mL of 70% EtOH. Following fixation, cells were pelleted by centrifuging for 5 min at 1400 rpm. Cells were resuspended in 1X PBS and pelleted again for 5 min at 1400 rpm. Residual PBS was decanted and cells were stained for 30min at room temperature with Propidium Iodide (Miltenyi Biotech, San Diego, CA) in the presence of RNase. Stained cells were analyzed for DNA content by fluorescence-activated cell sorting using a FACScan flow cytometer (Becton Dickinson). Percentages of cells in the G1, S, or G2/M phases of the cell cycle were quantified with FlowJo software. For each sample 20000 events were collected. Three biological replicates were performed, and representative data are shown.

#### ChIP-Sequencing

##### ChIP-seq reactions and antibodies

KCNR and LAN5 cells with or without ATRA treatment were subjected to Chromatin immunoprecipitation (ChIP) using ChIP-IT High Sensitivity kit (Active Motif) as per manufacturer’s protocol. Briefly, formaldehyde fixed cells (∼ 10 million cells) were flash frozen and stored in −80°C for later use. Nuclei were isolated and the chromatin was sheared in the range of 200-1000 base pairs using either an Active Motif EpiShear Probe Sonicator (Active Motif). KCNR cells were sonicated at 30% amplitude pulse for 20s on and 30s off for a total sonication “on” time of 14 min, LAN5 cells were sonicated at 25% amplitude pulse for 20s on and 30s off for a total sonication “on” time of 13 min. Sheared chromatin samples were immunoprecipitated overnight at 4°C with antibodies targeting H3K27ac (Active Motif, catalog #39133), H3K4me3 (Cell Signaling, catalog # 9751), H3K4me1 (Abcam, catalog # ab8895), and H3K9me3 (Active Motif, catalog # ab8898). DNA purifications were performed with the ChIP-IT High Sensitivity kit (Active Motif). To normalize ChIP-seq signal, we used Active Motif ChIP-seq spike in using Drosophila chromatin (Active Motif catalog # 53083) and an antibody against Drosophila specific histone variant H2Av (Active Motif, catalog #61686) according to the manufacturer’s instructions. ChIP samples were subjected to qPCR using ChIP-qPCR kit (Active Motif) as per manufacturers instruction. The primers are listed in Suppl. Table 10. ChIP-seq DNA libraries were prepared by Frederick National Laboratory for Cancer Research sequencing facility using Illumina TruSeq ChIP Library Prep Kit, after which DNA was size-selected with SPRIselect reagent kit (to obtain a 250-300 bp average insert fragment size). Libraries were multiplexed and sequenced using TruSeq ChIP Samples Prep Kit (75 cycles), cat. # IP-2-2-1012/1024 on an Illumina NextSeq500 machine. 35,000,000–45,000,000 reads were generated per sample.

##### ChIP-Seq Data Analysis

###### Data Processing

ChIP-seq data analysis was done as previously described, using similar pipelines and parameters ^60,61^. Briefly, ChIP-seq reads were aligned to reference human genome (hg19) using BWA ^62^ to build version GRCh37 (hg19) of the human genome. SAM files were converted to BAM files using SAMTools ^63^ and sorted, de-duplicated with picard and indexed using SAMTools. To visualize ChIP-seq tracks in IGV, BAM files were converted to tdf files using IGV tools (version 2.3.57). ChIP-Seq peaks were detected using MACS2 (version 2.1.0) ^64^. The default p value cut off for peak enrichment was set to 10^−5^ for all data sets. Peaks within 2,500bp to the nearest TSS were set as promoter proximal, while all other were considered distal. The distribution of peaks (as intronic, intergenic, exonic, etc.) was annotated using HOMER. ChIP-seq read density values were normalized per million mapped reads for a particular region. Gene ontology was performed using GREAT ^20^, using hg19 and the whole genome as the background.

###### Identification of enhancers and super-enhancers

Enhancers were detected using H3K27ac as an enhancer mark. Active enhancers were ranked according to increasing H3K27ac levels. To detect super-enhancers we used modified ROSE2 algorithm (https://bitbucket.org/young_computation/rose) ^65,66^ to take care of MYCN amplification and ranked the enhancers that were identified using MACS. The peaks were ranked according to the H3K27ac peak intensity (length × density) with input signal in the corresponding region subtracted. Peaks within 12.5 kb were stitched, however peaks within ± 2000 bases of an annotated promoter were excluded from stitching. The point where y=x was tangent to the curve formed by plotting the rank ordered stitched enhancer was selected as the threshold separating super-enhancers (SE) from typical enhancers (TE). Each enhancer was associated with the RefSeq genes that either overlapped with the enhancer, or whose transcriptional start sites (TSS) fell within 50kb of the enhancer boundaries. Differential peak calling between control and ATRA treated samples was performed using BEDTools v2.25.0 ^67^ in multicov mode to measure read counts, which were normalized per million mapped reads, and visualized using R package ggplot2 or NGS plot ^68^. A Pearson correlation coefficient (r) was calculated to measure the linear correlation between SE signal and mRNA expression of the downstream gene of the interest.

###### K-means clustering analysis

Clustering of SEs in ATRA treated KCNR cells was performed by first creating a master group of all called SEs from all timepoints and conditions, and then calculating the H3K27ac read count pileup in each SE over the treatment time-course. We removed SEs that did not change between treatment and control. Then, each region was normalized by Z-scoring the H3K27ac signal over all time points, followed by K means clustering using the R statistical package base function kmeans (x, centers = 5). The clustering separated SEs into groups that lost signal, or gained signal at different timepoints that we termed “waves” of SE activation.

###### RPMPR data normalization

For KCNR we normalized H3K27ac according to reads per million peak reads (RPMPR) by first calling peaks with MACS2, then calculating the total number of reads only within peaks and normalizing to million reads within peaks.

###### Linked Enhancer Activated Factor (LEAF) plots

To create LEAF plots, we used the edge output from COLTRON ^36^ that reports for each TF_a_ the list of other TFs_(a…z)_ where its motif is found in that TFs SE. Then, for each TF node we extracted the expression level from RNA-seq data paired to the same cell line, treatment condition and timepoint, and used expression to determine node rank and size. Then, arrows were drawn from edge connection data.

###### Hi-C and data processing

KCNR cells for Hi-C were prepared following the user guide of ARIMA GENOMICS Arima-HiC kit. Briefly, the cells were crosslinked and digested using Arima restriction enzyme cocktail. The ends were filled and marked with biotin and ligated. After DNA shearing and size selection, biotin-labeled ligation junctions were enriched using streptavidin beads. Libraries were prepared using Accel-NGS 2S Plus DNA library kit (Swift Biosciences), following the manufacturers protocol, which were then PCR-amplified. The resulting libraries were multiplexed and sequenced (150 bp pair end) with NovaSeq 6000 machine. Approximately 500 million reads were generated for each sample. The HiC-Pro (https://github.com/nservant/HiC-Pro) and Juicer pipeline (https://github.com/aidenlab/juicer) were used to process the Hi-C data, and a custom Arima-HiC-specific Juicer.sh command line script was integrated during this process. Juicebox was used to visualize and create the Hi-C maps.

###### ATAC-seq

ATAC-seq was performed as previously described ^69,70^. Approximately, 50,000 KCNR cells were pelleted and the Tn5 transposition reaction was performed with the Nextera kit (Illumina), according to the manufacturer’s protocol. ATAC libraries were sequenced on an Illumina NextSeq machine (paired-end 75-bp reads). The Fastq files were processed using Encode ATAC_DNase_pipelines (https://github.com/kundajelab/atac_dnase_pipelines) installed on the NIH biowulf cluster (https://hpc.nih.gov/apps/atac_dnase_pipelines.html).

###### Accession Numbers

The raw sequencing data and processed files were deposited in the GEO repository at the NCBI (GEO: GSE147408).

## Code availability statement

Code used to analyze the data is available here: https://qithub.com/CBIIT/ChlPseq and questions regarding its implementation can be directed to the corresponding authors.

## Reference

1 Maris, J. M. Recent advances in neuroblastoma. N Engl J Med 362, 2202–2211, doi: 10.1056/NEJMra0804577 (2010).

2 Guo, X., Chen, Q. R., Song, Y. K., Wei, J. S. & Khan, J. Exon array analysis reveals neuroblastoma tumors have distinct alternative splicing patterns according to stage and MYCN amplification status. BMC Med Genomics 4, 35, doi: 10.1186/1755-8794-4-35 (2011).

3 Maris, J. M. The biologic basis for neuroblastoma heterogeneity and risk stratification. Curr Opin Pediatr 17, 7–13, doi: 10.1097/01.mop.0000150631.60571.89 (2005).

4 Maris, J. M., Hogarty, M. D., Bagatell, R. & Cohn, S. L. Neuroblastoma. Lancet 369, 2106–2120, doi: 10.1016/S0140-6736(07)60983-0 (2007).

5 Boeva, V. et al. Heterogeneity of neuroblastoma cell identity defined by transcriptional circuitries. Nat Genet 49, 1408–1413, doi: 10.1038/ng.3921 (2017).

6 Durbin, A. D. et al. Selective gene dependencies in MYCN-amplified neuroblastoma include the core transcriptional regulatory circuitry. Nat Genet 50, 1240–1246, doi: 10.1038/s41588-018-0191-z (2018).

7 van Groningen, T. et al. Neuroblastoma is composed of two super-enhancer-associated differentiation states. Nat Genet 49, 1261–1266, doi: 10.1038/ng.3899 (2017).

8 Matthay, K. K. et al. Neuroblastoma. Nat Rev Dis Primers 2, 16078, doi: 10.1038/nrdp.2016.78 (2016).

9 Schulte, J. H. & Eggert, A. Neuroblastoma. Crit Rev Oncog 20, 245–270, doi: 10.1615/critrevoncog.2015014033 (2015).

10 Garraway, L. A. & Sellers, W. R. Lineage dependency and lineage-survival oncogenes in human cancer. Nat Rev Cancer 6, 593–602, doi: 10.1038/nrc1947 (2006).

11 Bradner, J. E., Hnisz, D. & Young, R. A. Transcriptional Addiction in Cancer. Cell 168, 629–643, doi: 10.1016/j.cell.2016.12.013 (2017).

12 Soldatov, R. et al. Spatiotemporal structure of cell fate decisions in murine neural crest. Science 364, doi: 10.1126/science.aas9536 (2019).

13 Kerosuo, L. et al. Enhanced expression of MycN/CIP2A drives neural crest toward a neural stem cell-like fate: Implications for priming of neuroblastoma. Proc Natl Acad Sci U S A 115, E7351–E7360, doi: 10.1073/pnas.1800039115 (2018).

14 Wakamatsu, Y., Watanabe, Y., Nakamura, H. & Kondoh, H. Regulation of the neural crest cell fate by N-myc: promotion of ventral migration and neuronal differentiation. Development 124, 1953–1962 (1997).

15 Cheung, N. K. & Dyer, M. A. Neuroblastoma: developmental biology, cancer genomics and immunotherapy. Nat Rev Cancer 13, 397–411, doi: 10.1038/nrc3526 (2013).

16 Wang, L. et al. ASCL1 is a MYCN- and LMO1-dependent member of the adrenergic neuroblastoma core regulatory circuitry. Nature Communications 10, 5622, doi: 10.1038/s41467-019-13515-5 (2019).

17 Gaetano, C., Matsumoto, K. & Thiele, C. J. Retinoic acid negatively regulates p34cdc2 expression during human neuroblastoma differentiation. Cell Growth Differ 2, 487–493 (1991).

18 Thiele, C. J., Reynolds, C. P. & Israel, M. A. Decreased expression of N-myc precedes retinoic acid-induced morphological differentiation of human neuroblastoma. Nature 313, 404–406, doi: 10.1038/313404a0 (1985).

19 Chipumuro, E. et al. CDK7 inhibition suppresses super-enhancer-linked oncogenic transcription in MYCN-driven cancer. Cell 159, 1126–1139, doi: 10.1016/j.cell.2014.10.024 (2014).

20 McLean, C. Y. et al. GREAT improves functional interpretation of cis-regulatory regions. Nat Biotechnol 28, 495–501, doi: 10.1038/nbt.1630 (2010).

21 Ritenour, L. E., Randall, M. P., Bosse, K. R. & Diskin, S. J. Genetic susceptibility to neuroblastoma: current knowledge and future directions. Cell Tissue Res 372, 287–307, doi: 10.1007/s00441-018-2820-3 (2018).

22 Maris, J. M. et al. Chromosome 6p22 locus associated with clinically aggressive neuroblastoma. The New England journal of medicine 358, 2585–2593, doi: 10.1056/NEJMoa0708698 (2008).

23 Capasso, M. et al. Common variations in BARD1 influence susceptibility to high-risk neuroblastoma. Nature genetics 41, 718–723, doi: 10.1038/ng.374 (2009).

24 Wang, K. et al. Integrative genomics identifies LMO1 as a neuroblastoma oncogene. Nature 469, 216–220, doi: 10.1038/nature09609 (2011).

25 Nguyen le, B. et al. Phenotype restricted genome-wide association study using a gene-centric approach identifies three low-risk neuroblastoma susceptibility Loci. PLoS Genet 7, e1002026, doi: 10.1371/journal.pgen.1002026 (2011).

26 Diskin, S. J. et al. Common variation at 6q16 within HACE1 and LIN28B influences susceptibility to neuroblastoma. Nature genetics 44, 1126–1130, doi: 10.1038/ng.2387 (2012).

27 Diskin, S. J. et al. Rare variants in TP53 and susceptibility to neuroblastoma. J Natl Cancer Inst 106, dju047, doi: 10.1093/jnci/dju047 (2014).

28 McDaniel, L. D. et al. Common variants upstream of MLF1 at 3q25 and within CPZ at 4p16 associated with neuroblastoma. PLoS Genet 13, e1006787, doi: 10.1371/journal.pgen.1006787 (2017).

29 Hungate, E. A. et al. Evaluation of Genetic Predisposition for MYCN-Amplified Neuroblastoma. J Natl Cancer Inst 109, doi: 10.1093/jnci/djx093 (2017).

30 Diskin, S. J. et al. Copy number variation at 1q21.1 associated with neuroblastoma. Nature 459, 987–991, doi: 10.1038/nature08035 (2009).

31 McDaniel, L. D. et al. Common variants upstream of MLF1 at 3q25 and within CPZ at 4p16 associated with neuroblastoma. PLoS genetics 13, e1006787 (2017).

32 Oldridge, D. A. et al. Genetic predisposition to neuroblastoma mediated by a LMO1 super-enhancer polymorphism. Nature 528, 418–421, doi: 10.1038/nature15540 (2015).

33 Decaesteker, B. et al. TBX2 is a neuroblastoma core regulatory circuitry component enhancing MYCN/FOXM1 reactivation of DREAM targets. Nat Commun 9, 4866, doi: 10.1038/s41467-018-06699-9 (2018).

34 Potzner, M. R. et al. Sequential requirement of Sox4 and Sox11 during development of the sympathetic nervous system. Development 137, 775–784, doi: 10.1242/dev.042101 (2010).

35 El Wakil, A., Francius, C., Wolff, A., Pleau-Varet, J. & Nardelli, J. The GATA2 transcription factor negatively regulates the proliferation of neuronal progenitors. Development 133, 2155–2165, doi: 10.1242/dev.02377 (2006).

36 Lin, C. Y. et al. Active medulloblastoma enhancers reveal subgroup-specific cellular origins. Nature 530, 57–62, doi: 10.1038/nature16546 (2016).

37 Kamachi, Y. & Kondoh, H. Sox proteins: regulators of cell fate specification and differentiation. Development 140, 4129–4144, doi: 10.1242/dev.091793 (2013).

38 Thakore, P. I. et al. Highly specific epigenome editing by CRISPR-Cas9 repressors for silencing of distal regulatory elements. Nat Methods 12, 1143–1149, doi: 10.1038/nmeth.3630 (2015).

39 Vermeulen, J. et al. Predicting outcomes for children with neuroblastoma using a multigene-expression signature: a retrospective SIOPEN/COG/GPOH study. The lancet oncology 10, 663–671 (2009).

40 Ohira, M. et al. Expression profiling using a tumor-specific cDNA microarray predicts the prognosis of intermediate risk neuroblastomas. Cancer cell 7, 337–350 (2005).

41 Gryder, B. E. et al. Histone hyperacetylation disrupts core gene regulatory architecture in rhabdomyosarcoma. Nat Genet 51, 1714–1722, doi: 10.1038/s41588-019-0534-4 (2019).

42 Dobin, A. et al. STAR: ultrafast universal RNA-seq aligner. Bioinformatics 29, 15–21, doi: 10.1093/bioinformatics/bts635 (2013).

43 Bergsland, M. et al. Sequentially acting Sox transcription factors in neural lineage development. Genes Dev 25, 2453–2464, doi: 10.1101/gad.176008.111 (2011).

44 Hoser, M. et al. Sox12 deletion in the mouse reveals nonreciprocal redundancy with the related Sox4 and Sox11 transcription factors. Mol Cell Biol 28, 4675–4687, doi: 10.1128/MCB.00338-08 (2008).

45 Chen, C., Jin, J., Lee, G. A., Silva, E. & Donoghue, M. Cross-species functional analyses reveal shared and separate roles for Sox11 in frog primary neurogenesis and mouse cortical neuronal differentiation. Biol Open 5, 409–417, doi: 10.1242/bio.015404 (2016).

46 Chen, C., Lee, G. A., Pourmorady, A., Sock, E. & Donoghue, M. J. Orchestration of Neuronal Differentiation and Progenitor Pool Expansion in the Developing Cortex by SoxC Genes. J Neurosci 35, 10629–10642, doi: 10.1523/JNEUROSCI.1663-15.2015 (2015).

47 Hur, W. et al. SOX4 overexpression regulates the p53-mediated apoptosis in hepatocellular carcinoma: clinical implication and functional analysis in vitro. Carcinogenesis 31, 1298–1307, doi: 10.1093/carcin/bgq072 (2010).

48 Pan, X. et al. Induction of SOX4 by DNA damage is critical for p53 stabilization and function. Proc Natl Acad Sci U S A 106, 3788–3793, doi: 10.1073/pnas.0810147106 (2009).

49 Zhu, S. et al. LMO1 Synergizes with MYCN to Promote Neuroblastoma Initiation and Metastasis. Cancer Cell 32, 310–323 e315, doi: 10.1016/j.ccell.2017.08.002 (2017).

50 Selmi, A. et al. TWIST1 is a direct transcriptional target of MYCN and MYC in neuroblastoma. Cancer Lett 357, 412–418, doi: 10.1016/j.canlet.2014.11.056 (2015).

51 Zeid, R. et al. Enhancer invasion shapes MYCN-dependent transcriptional amplification in neuroblastoma. Nat Genet 50, 515–523, doi: 10.1038/s41588-018-0044-9 (2018).

52 Lasorella, A. et al. Id2 is critical for cellular proliferation and is the oncogenic effector of N-myc in human neuroblastoma. Cancer Res 62, 301–306 (2002).

53 Woo, C. W. et al. Use of RNA interference to elucidate the effect of MYCN on cell cycle in neuroblastoma. Pediatr Blood Cancer 50, 208–212, doi: 10.1002/pbc.21195 (2008).

54 Dodonova, S. O., Zhu, F., Dienemann, C., Taipale, J. & Cramer, P. Nucleosome-bound SOX2 and SOX11 structures elucidate pioneer factor function. Nature 580, 669–672 (2020).

55 Gaetano, C., Matsumoto, K. & Thiele, C. J. In vitro activation of distinct molecular and cellular phenotypes after induction of differentiation in a human neuroblastoma cell line. Cancer Res 52, 4402–4407 (1992).

56 Bolger, A. M., Lohse, M. & Usadel, B. Trimmomatic: a flexible trimmer for Illumina sequence data. Bioinformatics 30, 2114–2120, doi: 10.1093/bioinformatics/btu170 (2014).

57 Love, M. I., Huber, W. & Anders, S. Moderated estimation of fold change and dispersion for RNA-seq data with DESeq2. Genome Biol 15, 550, doi: 10.1186/s13059-014-0550-8 (2014).

58 Mootha, V. K. et al. PGC-1alpha-responsive genes involved in oxidative phosphorylation are coordinately downregulated in human diabetes. Nat Genet 34, 267–273, doi: 10.1038/ng1180 (2003).

59 Subramanian, A. et al. Gene set enrichment analysis: a knowledge-based approach for interpreting genome-wide expression profiles. Proc Natl Acad Sci U S A 102, 15545–15550, doi: 10.1073/pnas.0506580102 (2005).

60 Gryder, B. E. et al. PAX3-FOXO1 Establishes Myogenic Super Enhancers and Confers BET Bromodomain Vulnerability. Cancer Discov 7, 884–899, doi: 10.1158/2159-8290.CD-16-1297 (2017).

61 Yohe, M. E. et al. MEK inhibition induces MYOG and remodels super-enhancers in RAS-driven rhabdomyosarcoma. Sci Transl Med 10, doi: 10.1126/scitranslmed.aan4470 (2018).

62 Li, H. & Durbin, R. Fast and accurate short read alignment with Burrows-Wheeler transform. Bioinformatics 25, 1754–1760, doi: 10.1093/bioinformatics/btp324 (2009).

63 Li, H. et al. The Sequence Alignment/Map format and SAMtools. Bioinformatics 25, 2078–2079, doi: 10.1093/bioinformatics/btp352 (2009).

64 Zhang, Y. et al. Model-based analysis of ChIP-Seq (MACS). Genome Biol 9, R137, doi: 10.1186/gb-2008-9-9-r137 (2008).

65 Loven, J. et al. Selective inhibition of tumor oncogenes by disruption of super-enhancers. Cell 153, 320–334, doi: 10.1016/j.cell.2013.03.036 (2013).

66 Whyte, W. A. et al. Master transcription factors and mediator establish super-enhancers at key cell identity genes. Cell 153, 307–319, doi: 10.1016/j.cell.2013.03.035 (2013).

67 Quinlan, A. R. & Hall, I. M. BEDTools: a flexible suite of utilities for comparing genomic features. Bioinformatics 26, 841–842, doi: 10.1093/bioinformatics/btq033 (2010).

68 Shen, L., Shao, N., Liu, X. & Nestler, E. ngs.plot: Quick mining and visualization of next-generation sequencing data by integrating genomic databases. BMC Genomics 15, 284, doi: 10.1186/1471-2164-15-284 (2014).

69 Corces, M. R. et al. An improved ATAC-seq protocol reduces background and enables interrogation of frozen tissues. Nat Methods 14, 959–962, doi: 10.1038/nmeth.4396 (2017).

70 Liu, Z. et al. CASZ1 induces skeletal muscle and rhabdomyosarcoma differentiation through a feed-forward loop with MYOD and MYOG. Nat Commun 11, 911, doi: 10.1038/s41467-020-14684-4 (2020).

