## Supplemental Figures for "Lineage specific transcription factor waves reprogram neuroblastoma from self-renewal to differentiation" (from another version of this preprint: version 1, 2020-07-24)

**Supplementary Figures**

**Supplementary Figure 1**

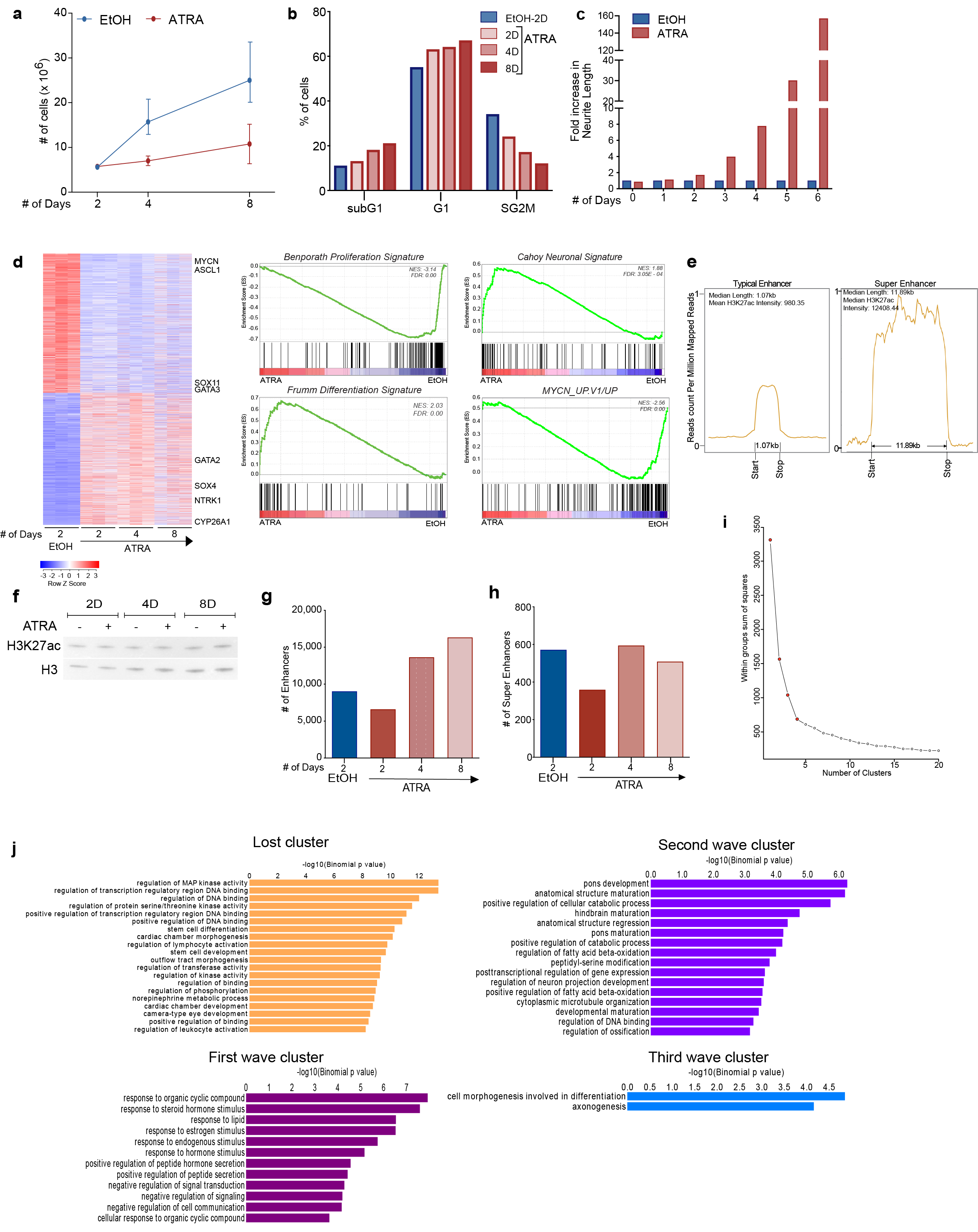

**Fig. S1 (Extension of Fig. 1): ATRA treatment leads to growth arrest and differentiation and changes in SE landscape of Neuroblastoma cells**

a. ATRA treatment leads to decrease in 50% decrease in cell number by 4 days and 90% by 8 days
b. ATRA induced growth arrest as measured by 20% increase in cells in G1 phase and 20% decrease in cells in G2/M phase by 8 days.
c. ATRA mediated differentiation was measured by 8-fold increase in neurite length by 4 days.
d. Left Panel: Heat map showing significant (FDR ≤ 0.01) differentially expressed genes after ATRA treatment. Right Panel: Gene set Enrichment analysis (GSEA) shows negative regulation of Benporath Proliferation and MYCN UP.V1/UP gene sets and positive enrichment of Frumm Differentiation and Cahoy neuronal gene set following ATRA treatment.
e. Metagene representations of the mean ChIP-seq signal across all typical enhancers and super-enhancer domains, identified in the self-renewing state. Metagenes are centered on the enhancer region, and the length of the enhancer reflects the difference in median lengths (1.07kb for typical enhancers, 11.9kb for super-enhancers). Additional 5 kb flanking each enhancer region is also shown.
f. Western blow analysis showing total levels of H3K27ac untreated or with 2, 4, and 8 days of ATRA treatment. H3 was used as loading control.
g. Total number of typical enhancers identified using H3K27ac Chip-seq under untreated or with 2, 4, and 8 days of ATRA treatment.
h. Total number of super-enhancers identified using H3K27ac Chip-seq under untreated or with 2, 4, and 8 days of ATRA treatment.
i. Elbow chart showing the SSE (sum of squared errors) after running k-means clustering for k going from 1 to 20. A clear elbow is seen at k = 4, indicating that 4 is the best number of clusters.
j. Gene ontology terms associated with genes found using GREAT analysis (great.stanford.edu) on H3K27ac constituent peaks within SEs in each cluster.

**Supplementary Figure 2**

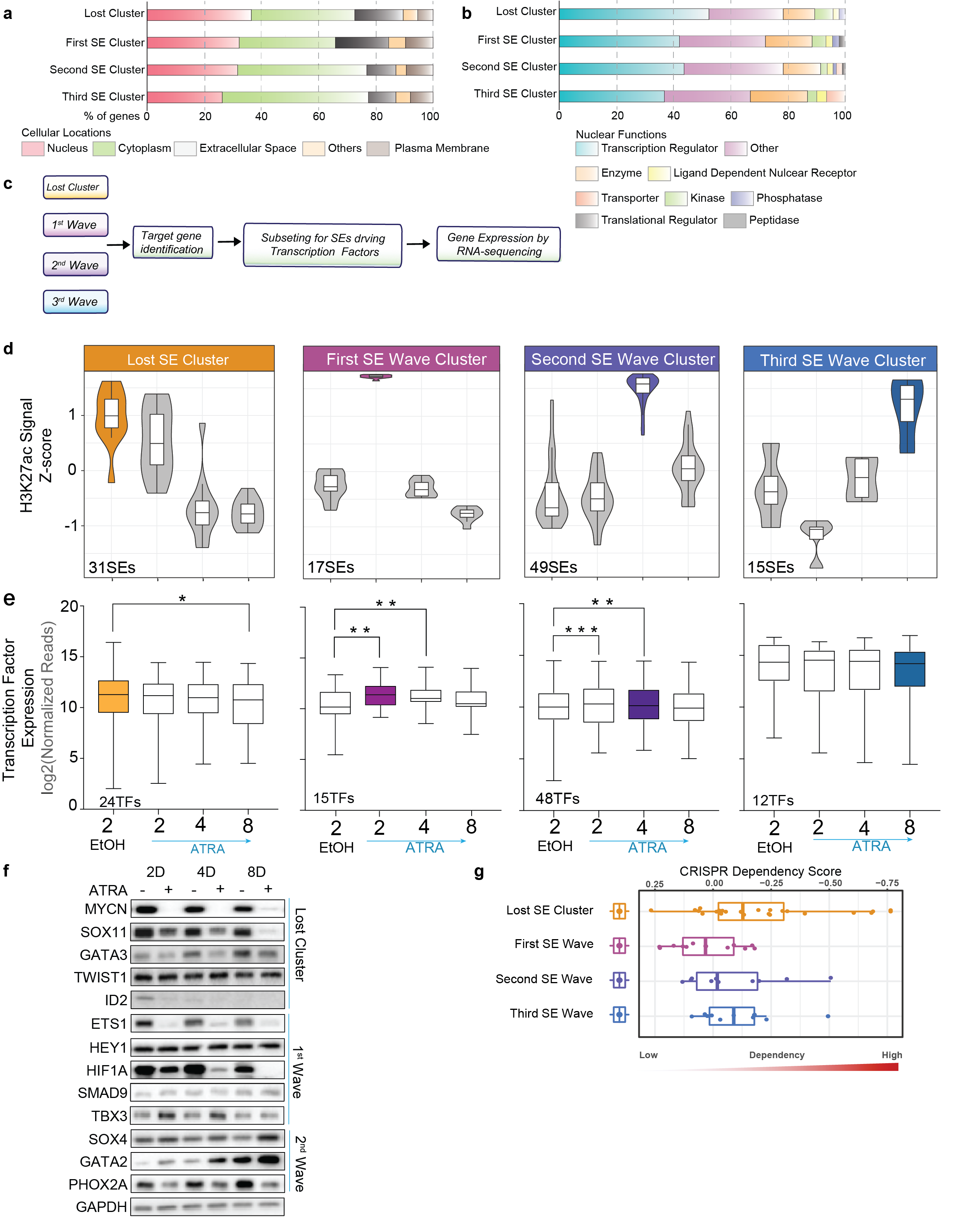

**Fig. S2 (Extension of Fig. 2): SE driven Transcription Factors drive proliferation and differentiation of KCNR cells.**

a. Stacked bar-chart showing the predicted cellular localization of the genes driven by the SEs in each cluster.
b. Stacked bar-chart showing the function of the SE cluster genes that are localized in the nucleus.
c. A schematic representation of the analysis steps following SEs identification in each cluster.
d. Violin plots of z-score of H3K27ac signal of SEs in each cluster. The lost cluster consists of 31SEs, the first wave cluster has 17SEs, the second wave cluster consists of 49 SEs, and the third cluster has 15SEs.
e. Bar graph showing expression of the TFs driven by the SEs in each cluster. The 31SEs in the lost cluster drives 24TFs where the loss in SE signal lead to significant decrease in the expression of the corresponding TFs after 8 days of ATRA treatment. Similarly gain in SE signal after 2 days and 4 days of ATRA treatment lead to significant gain in the expression of the TFs. However, no change in expression of SE driven TFs was observed for the third cluster.
f. Western blot analysis of selected TFs of 3 clusters after 2,4 and 8 days of vehicle or ATRA treatment.
g. Bar graph showing dependency of neuroblastoma cell line on all the TFs in each cluster for cell growth and proliferation. NB cells are most preferentially dependent on lost cluster TFs for growth and proliferation compared to TFs in other clusters. Data mined from Project Achilles genome wide CRISPR-Cas9 screen.

**Supplementary Figure 3**

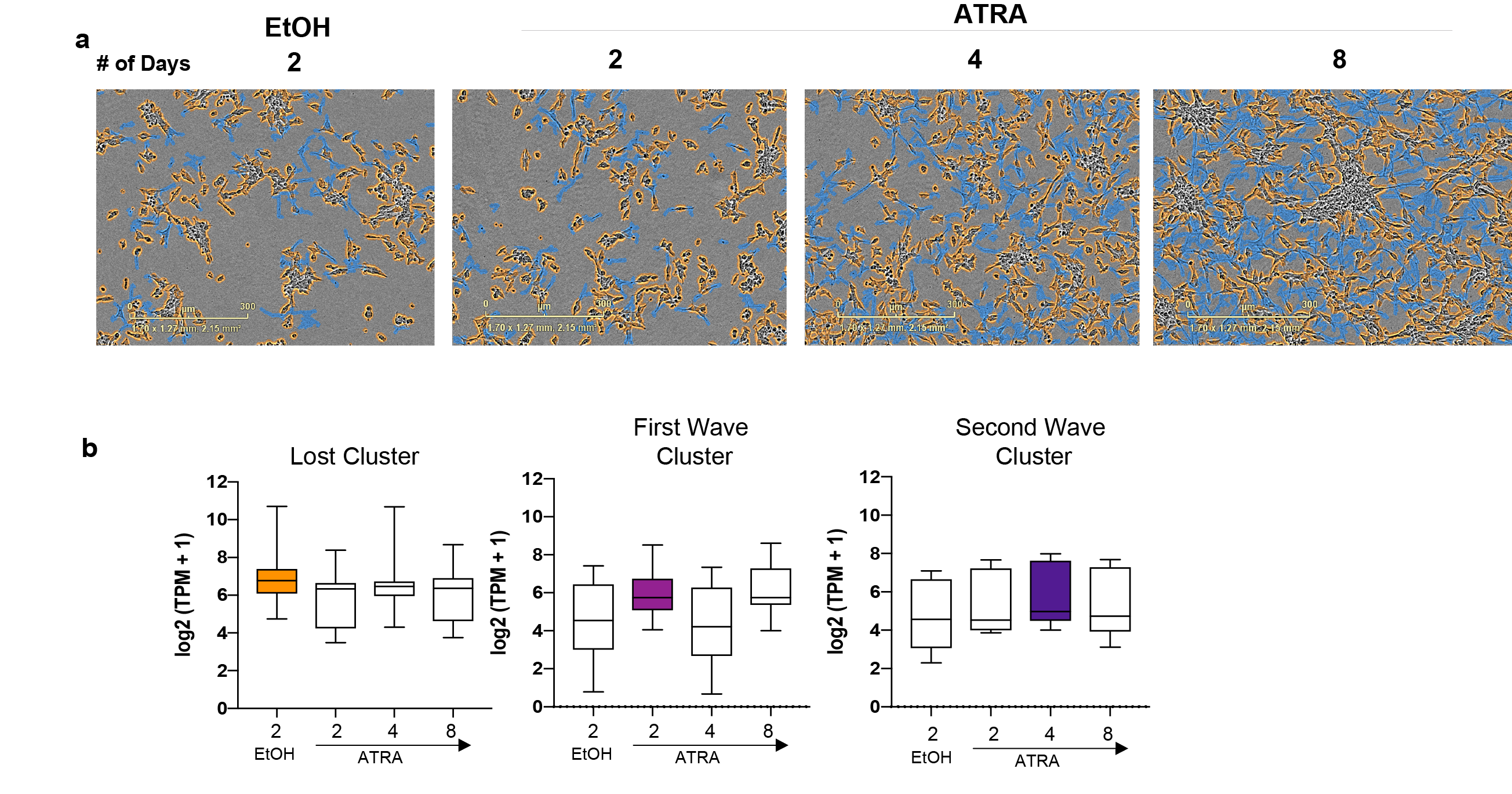

**Fig. S3 (Extension of Fig. 2): ATRA treatment of LAN5 cells drives differentiation and regulates expression of TFs.**

a. Representative images of LAN5 cells showing time dependent effect of 5µM ATRA treatment on cell morphology. The cell body is highlighted in yellow and the neurites are highlighted in blue.
b. Bar graph showing expression of the TFs driven by the SEs in each cluster (common with KCNR analyzed in Fig. 2C). Significant decrease in the expression of the TFs in lost cluster was observed 2, 4 and 8 days of ATRA treatment in LAN5 cells. Similarly, significant gain in expression of the TFs in first and second cluster was also observed in LAN5 cells after ATRA treatment.

**Supplementary Figure 4**

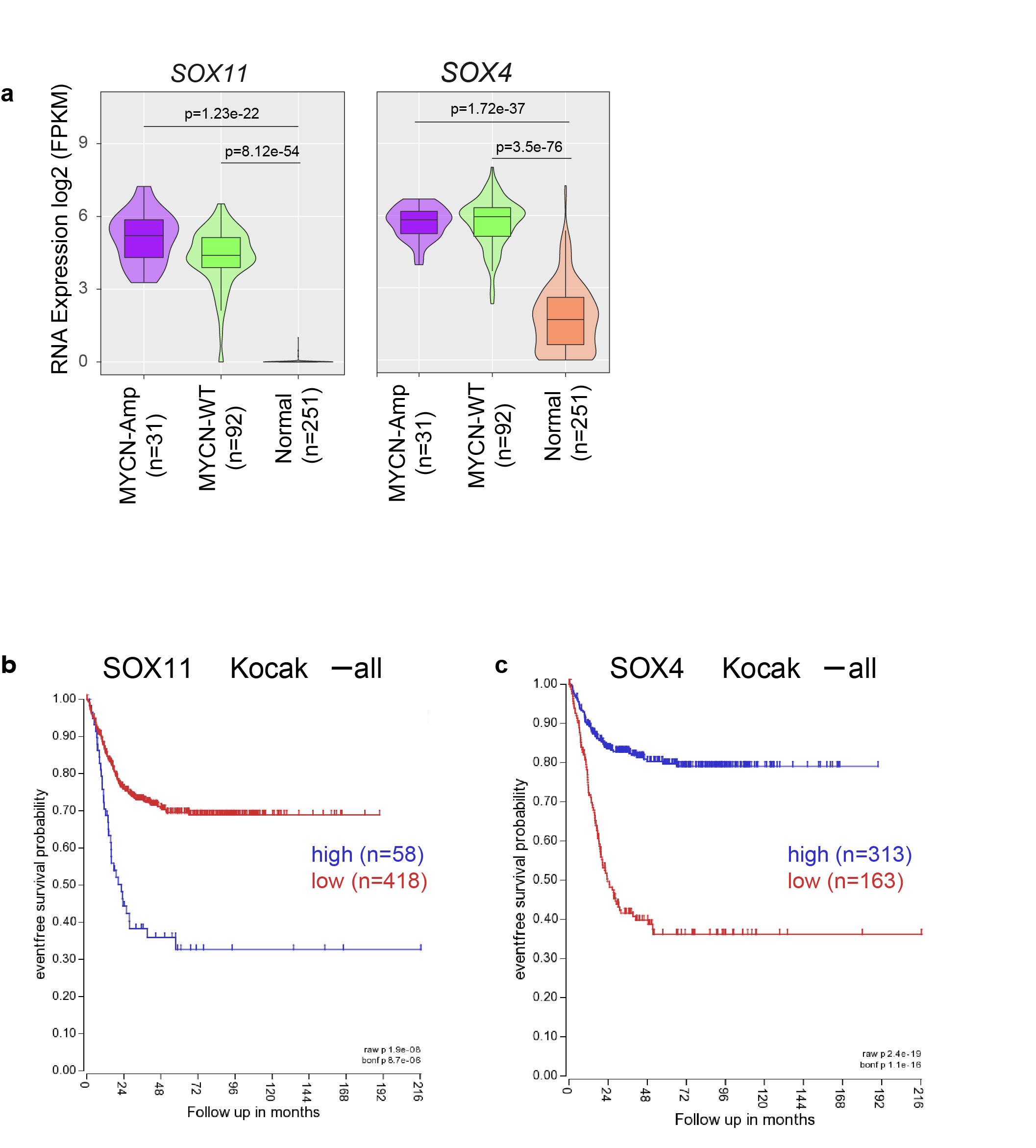

**Fig. S4 (Extension of Fig. 3): SOX11 and SOX4 are expressed in NB cells and associated with prognosis in NB Tumors**

A. Violin plots showing gene expression of SOX11 and SOX4 in MYCN-Amp neuroblastoma tumors, MYCN-WT neuroblastoma tumors, and normal tissues.
B. Kaplan-Meier plots based on the expression of SOX11 (left panel) and SOX4 (right panel) in tumors from NB patients at all stages (R2 database: Kocak dataset).

**Supplementary Figure 5**

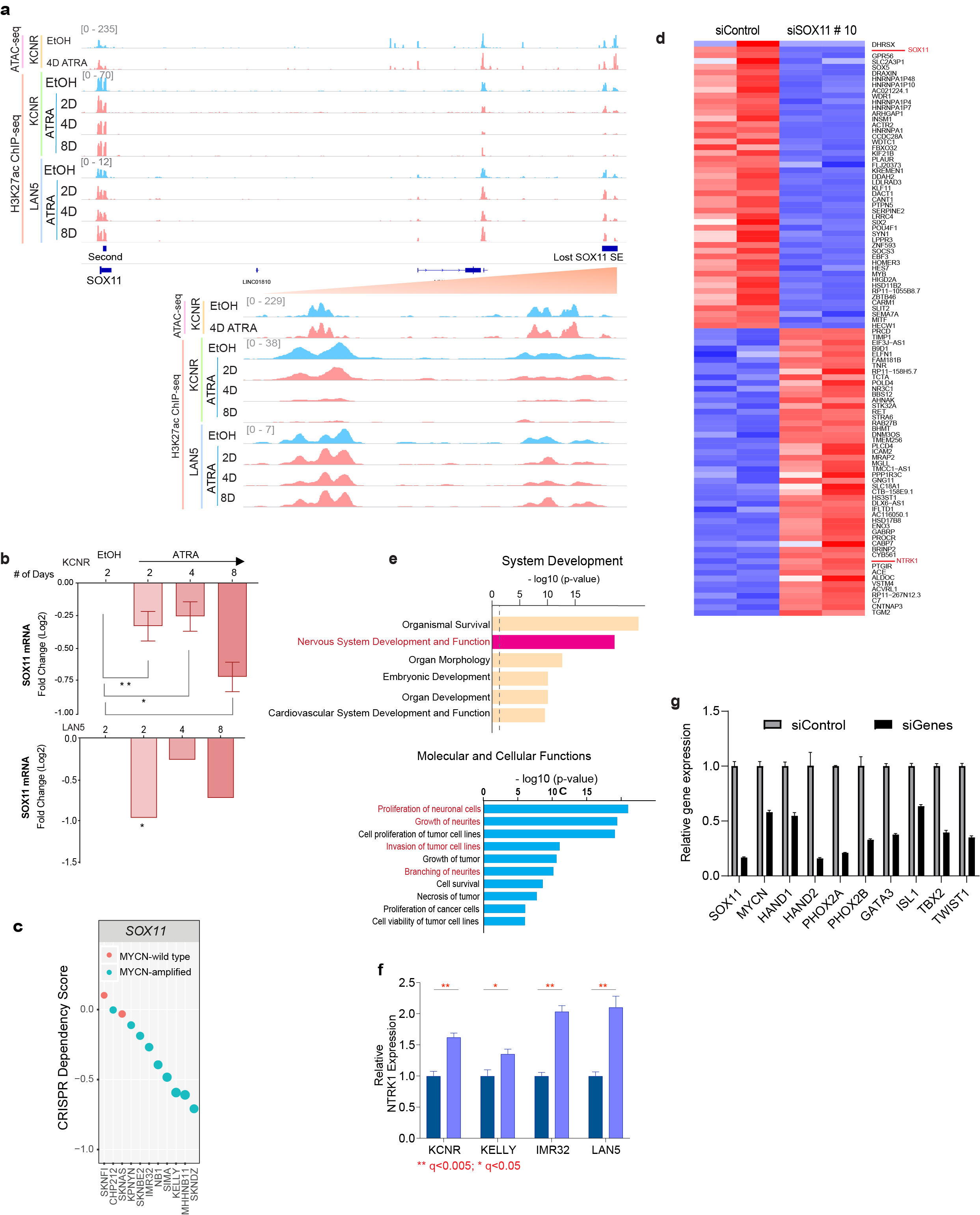

**Fig. S5 (Extension of Fig. 4 and 5): SOX11 expression is regulated by SEs in NB cells**

a. Representative ChIP-Seq tracks for H3K27ac in EtOH (blue) and ATRA treated cells (pink), showing loss of H3K27ac peaks at SOX11 SE in LAN5 cells. ChIP-seq read densities (y axis) are normalized and reported as reference-adjusted reads per million mapped reads (RRPM) in each sample. Super-enhancers are noted as purple bars.
b. Expression of SOX11 as determined by RNA-seq in KCNR and LAN5 cells in the presence and absence of ATRA. Log 0.75-fold increase in SOX11 expression was observed post ATRA treatment in both cell lines. Data are means ± SD for three replicates; P value generated from unpaired t test with Welch’s correction.
c. Dot plot showing the dependency of individual NB cell line on SOX11 for growth and proliferation. Data mined from Project Achilles genome wide CRISPR-Cas9 screen.
d. Heatmap showing top 50 upregulated and 50 downregulated genes ranked by statistical significance 48hrs post SOX11 inhibition in KCNR cells. Data are presented as normalized expression values of two biological replicates based on DESeq2 software analysis and p-adj <0.001. The color key represents the normalized expression values: blue (low) to red (high).
e. Most significant system development pathways regulated by SOX11 target genes (upper panel). The top molecular functions decreased by SOX11 silencing (lower panel) as defined by IPA based on DESeq2 differential gene expression analysis.
f. RT-qPCR analysis showing significant increase in NTRK1 relative mRNA levels after SOX11 silencing in NB cells.
g. RT-qPCR analysis showing decrease in gene expression after silencing of respective TFs in NB cells.
